# WMH-DualTasker: A weakly-supervised deep learning model for automated white matter hyperintensities segmentation and visual rating prediction

**DOI:** 10.1101/2024.11.12.623137

**Authors:** Yilei Wu, Zijian Dong, Hongwei Bran Li, Yao Feng Chong, Fang Ji, Joanna Su Xian Chong, Nathanael Ren Jie Tang, Saima Hilal, Huazhu Fu, Christopher Li-Hsian Chen, Juan Helen Zhou, the Alzheimer’s Disease Neuroimaging Initiative

## Abstract

**Background:** White matter hyperintensities (WMH) are neuroimaging markers linked to an elevated risk of cognitive decline. WMH severity is typically assessed via visual rating scales and through volumetric segmentation. While visual rating scales are commonly used in clinical practice, they offer limited descriptive power. In contrast, supervised volumetric segmentation requires manually annotated masks, which is labor-intensive and challenging to scale for large studies. Therefore, our goal was to develop an automated deep learning model that can provide accurate and holistic quantification of WMH severity with minimal supervision.

**Methods:** We developed WMH-DualTasker, a deep learning model that simultaneously performs voxel-wise segmentation and visual rating score prediction. The model employs self-supervised learning with transformation-invariant consistency constraints, using WMH visual ratings (ARWMC scale, range 0-30) from clinical settings as the sole supervisory signal. Additionally, we assessed its clinical utility by applying it to identify individuals with mild cognitive impairment (MCI) and to predict dementia conversion.

**Findings:** The volumetric quantification performance of WMH-DualTasker was either superior to or on par with existing supervised methods, as demonstrated on the MICCAI-WMH dataset (N=60, Dice=0.602) and the SINGER dataset (N=64, Dice=0.608). Furthermore, the model exhibited strong agreement with clinical visual rating scales on an external dataset (SINGER, MAE=1.880, K=0.77). Importantly, WMH severity metrics derived from WMH-DualTasker improved predictive performance beyond conventional clinical features for MCI classification (AUC=0.718, p<0.001), MCI conversion prediction (AUC=0.652, p<0.001) using the ADNI dataset.

**Interpretations:** WMH-DualTasker substantially reduces the reliance on labor-intensive manual annotations, facilitating more efficient and scalable quantification of WMH severity in large-scale population studies. This innovative approach has the potential to advance preventive and precision medicine by enhancing the assessment and management of vascular cognitive impairment associated with WMH.

Code and model weights are publicly available at https://github.com/hzlab/WMH-DualTasker.

## 1 Introduction

White matter hyperintensities (WMH) are neuroimaging markers that are usually indicative of cerebral small vascular disease — a condition characterized by damage to the small blood vessels in the brain. Using a T2-weighted fluid-attenuated inversion recovery (FLAIR) sequence on magnetic resonance imaging (MRI) [1, 2], these hyperintensities manifest as areas of abnormal heightened signal intensity in the white matter. The presence of WMH carries significant clinical implications: studies have demonstrated their association with a myriad of neurological complications, including accelerated cognitive decline, a heightened risk of dementia, and an increased probability of stroke [3–5]. These links underscore WMH crucial influence on cognition and cerebrovascular disease.

There are two main ways to quantify WMH: voxel-wise segmentation and visual rating (Figure 1). Segmentation provides a volumetric assessment and an objective measure of WMH volume. While volumetric measurements provide greater sensitivity in detecting differences between patient groups [58], they are time-consuming and technically challenging, making them more commonly used in research settings than in clinical practice. In contrast, visual rating is a more common quantification method in clinical settings, where trained raters assess WMH based on standardized scales [2, 6–8]. Trained raters, such as radiologists and neurologists, evaluate and grade the severity and distribution of WMH based on their appearance in the scans. Several visual rating scales have been developed and validated, including the Fazekas scale [61] and the Age-Related White Matter Changes (ARWMC) scale [7]. ARWMC grades WMH severity by assigning scores from 0 to 3 in five bilateral brain regions (frontal lobe, parietal-occipital lobe, temporal lobe, infratentorial region, and basal ganglia), resulting in a global score ranging from 0 to 30, where a higher score indicates increased severity. However, there are two main limitations: First, the subjective nature of visual ratings can introduce variability among different raters. Second, visual rating scales show ceiling effects and poor discrimination of WMH severity at higher lesion volumes, as demonstrated by the LADIS study [58]. The LADIS study also found that volumetric measurements provide greater sensitivity in detecting differences between patient groups, particularly for memory symptoms. These complementary strengths and limitations suggest that an ideal approach would combine both methods: visual ratings for efficient clinical assessment and volumetric measurements for precise quantification. This observation motivates our approach of simultaneously generating both measurements.

**Figure 1:**
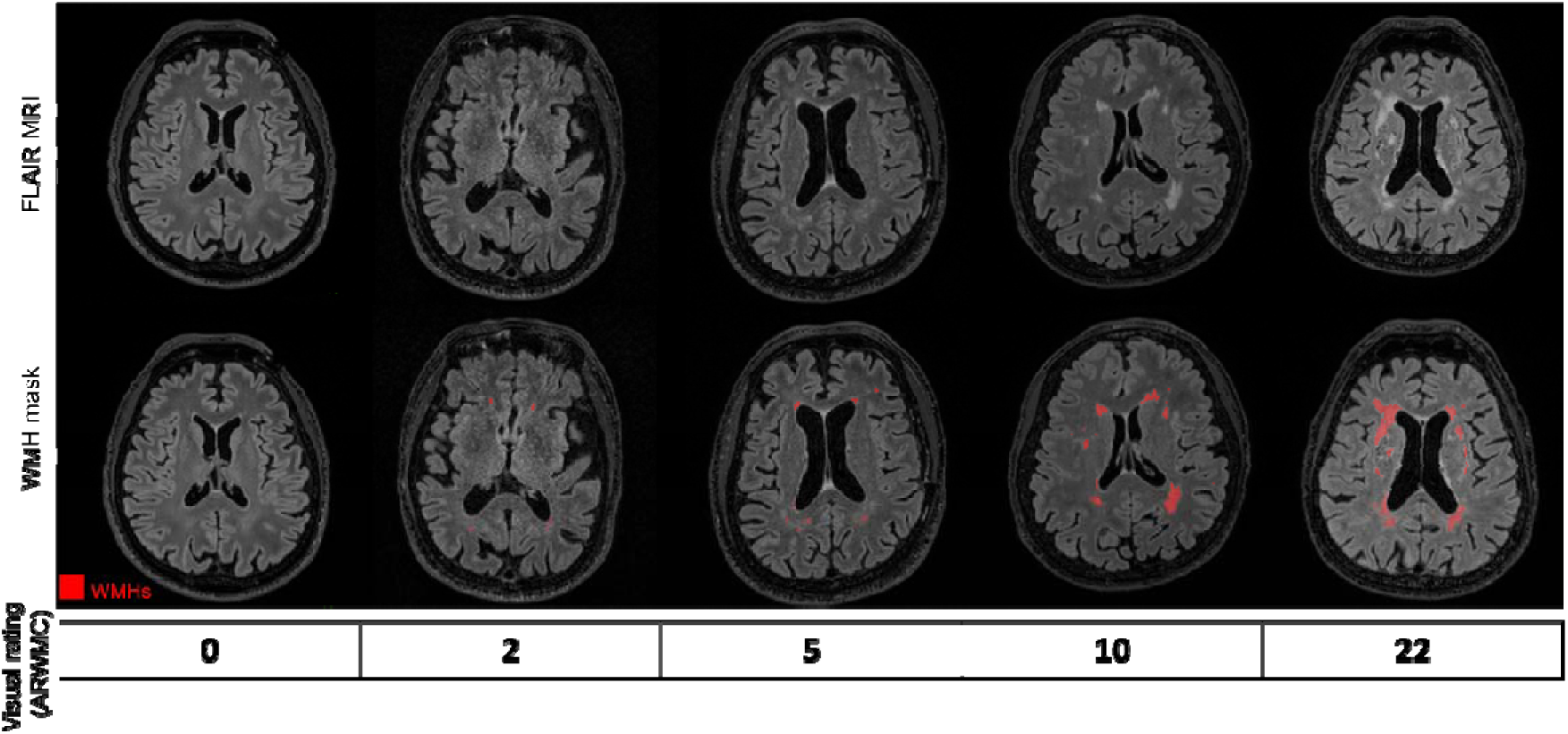
Illustration of WMH across varying loads. This figure presents a series of axial brain MRI scans from subjects, showcasing a gradient of WMH from minimal to pronounced, organized from left to right. In the top row, WMH is shown as high-intensity signals on T2-FLAIR MRI scans, contrasting sharply with the surrounding brain tissue and visually pinpointing the regions impacted by WMH. The bottom row complements this with WMH segmentation masks, rendered in striking red, to provide a precise visual quantification of the WMH load. We observe that WMH typically manifests as irregularly shaped, hyperintense lesions located in both periventricular and deep white matter regions on MRI scans. The final row features the ARWMC visual rating scale, introducing a standardized, qualitative approach to assess WMH severity. MRI: Magnetic Resonance Imaging. WMH: white matter hyperintensities. ARWMC: Age-Related White Matter Changes.

In the early era of automated WMH segmentation research, methodologies primarily rely on conventional machine learning techniques [9–11], including but not limited to random forest and logistic regression-based frameworks. Conventional machine learning methods relies on hand-designed feature extraction pipeline. Whereas, deep-learning method, with convolutional operators adjust feature automatically based on the data that has been provided during the training phase. Recent studies [12–14] highlight the preliminary successes of deep learning-based models for this task.

However, despite these advances, one outstanding challenge that remains is the dependence of these models on manually labelled WMH masks for effective training. The labelling process is labour-intensive; typically, a domain expert, such as a radiologist, dedicates between 0.5 to 2 hours to the meticulous labelling of a single MRI volume. This requirement for specialized expertise and the associated time commitment accentuates the difficulty in collecting large-scale manually labelled WMH datasets. Without a large number of such datasets, it is unlikely that the data will be sufficiently expansive and comprehensive to train a generalizable and performant segmentation model.

In this paper, we introduce WMH-DualTasker, a weakly supervised deep learning-based approach for simultaneous WMH segmentation and visual rating prediction. WMH-DualTasker predicts both the ARWMC score and the WMH mask using only visual rating scales for supervision, thus overcoming the challenge of acquiring manual WMH segmentation masks. The model leverages self-supervised, transformation-invariant consistency constraints to enhance WMH detection sensitivity while using clinical visual ratings as the sole supervisory signal. To the best of our knowledge, our work is the first to present a 3D weakly supervised segmentation framework guided by image-level weak supervision, distinct from methods guided by an object bounding box or partial annotations. Through evaluation on three distinct datasets, our framework demonstrates performance comparable to supervised methods [9–11] and shows high agreement with radiologist ratings. Furthermore, application to the Alzheimer’s Disease Neuroimaging Initiative (ADNI) dataset [15] demonstrates that WMH-DualTasker’s outputs can enhance the accuracy of distinguishing between severe and mild stages of cognitive impairment. This underscores WMH-DualTasker’s potential not only as a proficient tool for WMH identification but also as a clinically relevant aid in distinguishing between different stages of cognitive impairment.

## 2 Methods

### 2.1 Study design and datasets

Throughout our experiment, we used four datasets that span a range of ages from middle-aged to elderly, as well as multiple sites and ethnicities. Figure 2 illustrated our study design and configuration among the four datasets and the type of annotation they have. We described each dataset in the following part of this section and in Table 1, and more demographic information can be found in Tables S2-S5 in the supplementary material.

**Figure 2.**
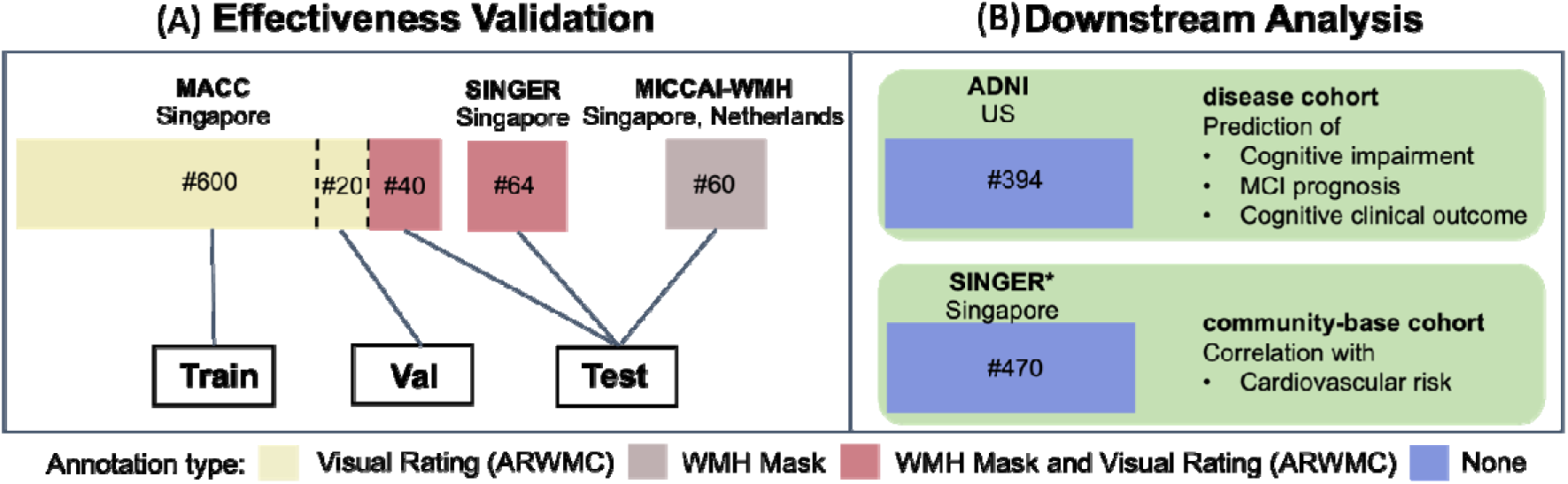
Comprehensive dataset utilization and downstream analysis framework for WMH-DualTasker. The schematic illustrates our multi-dataset approach and analytical pipeline. (A) For model development and evaluation, we utilized three primary datasets (MACC, SINGER, and MICCAI-WMH). The MACC dataset is divided into training, validation, and testing subsets, while SINGER and MICCAI-WMH are used as external test sets. Color-coding indicates the type of annotation available: yellow for visual ratings only, red for both visual ratings and manual segmentation masks, and grey for manual segmentation masks only. Subsequently, (B) We validate the utility of WMH-DualTask by performing our downstream analysis, demonstrating the application of WMH-DualTasker to two distinct cohorts: the ADNI dataset (a disease-specific cohort) and the SINGER dataset (a community-based cohort). These analyses encompass a range of clinically relevant tasks, including cognitive impairment classification, dementia conversion prediction, and assessment of dementia risk, showcasing the model’s versatility and potential clinical utility across diverse population settings.

**Table 1.** The overview of the scanner and image details in this study.

| **Dataset** | **Usage** | **Site** | **Cohort** | **Scanner** | **Sequence** | **Voxel resolution (mm<sup>3</sup>)** | **Number of subjects / cases with mask / cases with visual rating** |
| --- | --- | --- | --- | --- | --- | --- | --- |
| **MACC** | Train / Validation | Singapore | SVD | 3T Siemens Trio Tim | T2-FLAIR | 1.0x1.0x3.0 | 660 / 60 / 660 |
| **SINGER** | Validation | Singapore | SVD | 3T Siemens Trio Tim | T2-FLAIR | 1.0x1.0x1.0 | 60 / 60 / 60 |
| **SINGER\*** | Downstream Analysis | Singapore | SVD | 3T Siemens Trio Tim | T2-FLAIR | 1.0x1.0x1.0 | 470 / 0 / 0 |
| **MICCAI-UMC Utrecht** | Validation | Singapore | SVD | 3T Phillips Achieva | T2-FLAIR | 1.0x1.0x3.0 | 20 / 20 / 20 |
| **MICCAI-NUHS** | Validation | Netherlands | SVD | 3T Siemens Trio Tim | T2-FLAIR | 1.0x1.0x3.0 | 20 / 20 / 20 |
| **MICCAI-VU Amsterdam** | Validation | Netherlands | SVD | 3T GE Signa HDxT | T2-FLAIR | 1.0x1.0x1.2 | 20 / 20 / 20 |
| **ADNI** | Downstream Analysis | US | SVD, Control | 3T Siemens Trio Tim<br>3T Siemens Skyra<br>3T Philips Ingenia<br>1.5T Siemens Avanto | T2-FLAIR | 1.0x1.0x1.2 | 394 / 0 / 0 |

Note. SVD = Small Vessel Disease; T2-FLAIR = T2-weighted Fluid-Attenuated Inversion Recovery.

#### 2.1.1 MACC dataset

A total of 660 T2-FLAIR MRI images from the Memory, Ageing and Cognition Centre (MACC) dataset were used. This MACC dataset was part of an ongoing prospective memory clinic study. Participants with no cognitive impairment (NCI), cognitive impairment with no dementia (CIND), Alzheimer’s disease (AD) dementia and vascular dementia (VaD) were recruited from the National University Hospital of Singapore and Saint Luke’s Hospital [72–74]. Detailed demographic information used in our own analysis can be found in Supplementary Table S2. All images were quality-checked, and ARWMC ratings were performed by a team of experienced neuroradiologists, each with over six years of clinical neuroimaging experience. The ratings were then discussed with neurologists, psychologists, and research personnel during weekly consensus meetings [71]. For the MACC-Test dataset, WMH masks were generated through manual annotations of FLAIR images using ITK-SNAP (www.itksnap.org) and served as the reference standard for evaluating segmentation performance. The annotation process followed the STRIVE criteria [1,2] for standardized neuroimaging features of small vessel disease, with WMH defined as white matter hyperintensities on T2-weighted FLAIR images. To reduce noise artifacts, a minimum cluster size of 3 connected voxels was established. Each image was independently annotated by two experienced (over six year of experience) radiologists, with a third radiologist (over ten year of experience) reviewing cases where consensus was not reached. Overall, we had three splits: MACC-Train (n=600), MACC-Val (n=20), and MACC-Test (n=40) (see Figure 2). To prevent information leakage, we used MACC-Val for selecting hyper-parameters, and MACC-Test was hidden from the model development. It is worth mentioning that our model was trained on MACC-Train with image-level weak signals **only**, and the best model was selected based on the best ARWMC prediction the MACC-Val.

#### 2.1.2 MICCAI-WMH and SINGER dataset

We utilized a total of 60 images from the MICCAI-WMH challenge [16] and 64 images from the Singapore Geriatric Intervention Study (SINGER) [17] to examine the performance of the WMH-DualTasker across different datasets. The MICCAI-WMH was a public multi-site dataset from an open challenge, MRI scans with WMH masks that were manually annotated and cross-verified by two independent annotations from radiologists with over ten years of experience. The SINGER study was designed to assess the efficacy and safety of multidomain lifestyle interventions aimed at delaying cognitive decline in older adults at risk for dementia with specific risk factors for dementia. The SINGER study is a 2-year randomized controlled trial (RCT). 1200 older adults at risk of cognitive decline and dementia, determined using a Cardiovascular Risk Factors, Aging, and Incidence of Dementia (CAIDE) dementia risk score, will be recruited from community dwelling and clinical cohorts of Singaporean older adults from August 2021 [17]. For our analysis, we leveraged MRI scans from SINGER, accompanied by manually annotated WMH masks and visual ratings performed by radiologists using the ARWMC scale (the process is similar to the MACC). Complete demographic characteristics were provided in Supplementary Table S3. We evaluated our model using the MICCAI-WMH and SINGER as external testing datasets to evaluate the generalizability of the WMH-DualTasker for both WMH segmentation and regression.

#### 2.1.3 ADNI dataset

We employed the Alzheimer’s Disease Neuroimaging Initiative (ADNI) dataset to assess the clinical significance of the WMH-DualTasker model. Data used in the preparation of this article were obtained from the Alzheimer’s Disease Neuroimaging Initiative (ADNI) database (adni.loni.usc.edu). The ADNI was launched in 2003 as a public-private partnership, led by Principal Investigator Michael W. Weiner, MD. The primary goal of ADNI has been to test whether serial MRI, positron emission tomography (PET), other biological markers, and clinical and neuropsychological assessment can be combined to measure the progression of mild cognitive impairment (MCI) and early Alzheimer’s disease (AD). For up-to-date information, see www.adni-info.org. We selected 394 ADNI subjects aged 65-85 years who had completed cognitive evaluations and MRI scans at their baseline visits (Figure 2). According to their baseline diagnoses, these subjects comprised 152 Cognitively Normal (CN), 149 Mild Cognitive Impairment (MCI), and 93 Alzheimer’s Disease (AD) participants. Detailed demographic information for CN and MCI subjects can be found in Supplementary Table S4, and information about MCI prognosis subjects is available in Supplementary Table S5. Each subject’s MRI scan included FLAIR images was acquired in the axial plane.

#### 2.1.4 Ethics

The MACC study the ethics approval from NHG DOMAIN SPECIFIC REVIEW BOARD (DSRB) APPROVAL with a reference number of 2015/00406 (approval period: 01 November 2017 to 31 March 2020). The NHG DSRB operates in accordance to the ICH GCP and all applicable laws and regulations. Subjects were recruited into the study from the NUHS memory clinics and NUHS MACC research cohorts. Considering the retrospective nature of this study, the informed consent was waived.

The SINGER study obtained ethics approval from NUS-IRB REVIEW BOARD APPROVAL with a reference number of 2020/398 (approval period: 28 Aug 2020 to 31 May 2022). This is human biomedical research that is regulated by the Human Biomedical Research Act (HBRA) and researchers are required by law to comply with all the relevant regulatory requirements of the HBRA.

WMH-MICCAI dataset was available through the website (https://wmh.isi.uu.nl/). The MACC and SINGER datasets can be accessed through our data catalogue (https://macc-sg.notion.site/Data-Catalogue-122e6ba0007d80688d11fa10b6fcbd01). Researchers interested in accessing these datasets can submit their request through our online form (https://nus.syd1.qualtrics.com/jfe/form/SV_5psiKZDdeCoSQL4).

#### 2.1.5 Pre-processing

We implemented a standardized pre-preprocessing pipeline across all datasets. Given that our training datasets (MACC) was acquired at 1.00×1.00×3.00mm³ resolution, we resampled all validation images to match this resolution using trilinear interpolation. To determine appropriate cropping dimensions while ensuring complete brain coverage, we employed a two-step quality control process: (1) brain extraction used FSL-BET followed by manual visual verification, and (2) cropping size was determined based on the extracted binary brain masks to prevent any tissue loss. Based on this analysis, we established standardized dimensions of 256×256×64 for all preprocessed images. Finally, we applied z-score transformation to standardize the intensity distribution across all samples.

### 2.2 WMH-DualTasker - automatic segmentation and visual rating of WMH

This section introduced the formulation of WMH-DualTasker, which aimed to provide a holistic solution for automated WMH analysis by simultaneously accomplishing two tasks: WMH segmentation and ARWMC prediction. It was trained in an end-to-end manner, using only the ARWMC score as the supervisory signal. We began with the challenge of identifying White Matter Hyperintensities (WMH) using image-level weak labels. Then, we described how self-supervised equivalent regularization loss enhances this process, and how integrating a hyperintense map further optimized it by leveraging the prior characteristics of WMH. Finally, in Section 2.2.3 the inference strategy employed was introduced. WMH-DualTasker utilized a 3D model and takes MRI volumes as its input (see Figure 3). More details about the model structure can be found in Figure S1 in the supplementary material.

**Figure 3.**
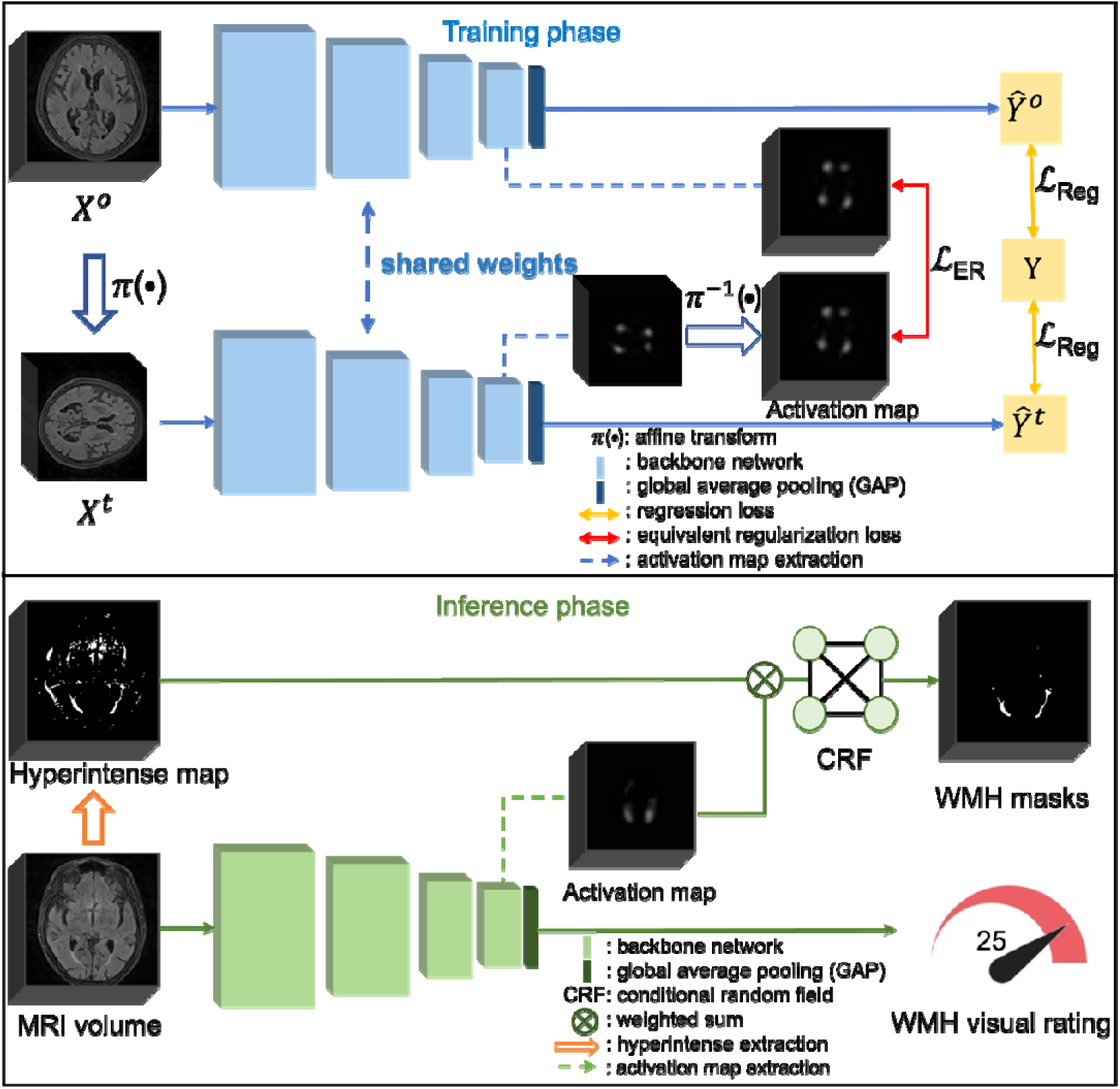
Schematic diagram of the WMH-DualTasker training and inference processes. The top portion of the diagram depicts the training phase, supervised by WMH visual rating and self-supervised equivalent regularization loss. The bottom part illustrates the inference process, where the enhanced CAM is integrated with the Hyperintense Map (HM) to produce the final WMH segmentation. CAM: Class Activation Map.

#### 2.2.1 Weakly supervised WMH segmentation through optimized CAM with self-supervised transformation equivalent regularization loss

Our approach to weakly supervised WMH segmentation centered on optimizing the Class Activation Map (CAM [19]) technique, a powerful tool for visualizing influential regions in image classification tasks. It worked by generating a salient map that highlights the parts of an image most relevant to a particular class label. In this work, we extracted a CAM that highlights WMH by training a 3D convolutional neural network to predict ARWMC given a brain MRI volume. We adopted the 3D light-weight structure (Simple Fully Convolutional Network, SFCN) [20] as the backbone network in our method (unless stated otherwise) due to its lower memory footprint. We described the detailed network configuration in Figure S1.

Due to the weak supervisory signal in the classification task compared to the segmentation task, the visualization of CAM was usually blurry and thus cannot accurately localize white matter abnormalities. In addition, as WMH frequently appeared in periventricular areas [4], we observed that the CAM simply highlights the centre of the brain. This motivated us to improve the localization capabilities of CAM from two perspectives. First, we refined its resolution by eliminating the max-pooling layers from standard 3D backbone networks, thereby obtaining feature maps with a higher spatial resolution. Specifically, our feature maps had a resolution of 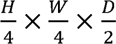 where H, W, D indicated the height, width, and depth of the input volume. Second, we introduced a regularization loss to address the limited localization capability of CAM from the regression task. Using CAM as an initial seed would result in poor WMH segmentation quality. To overcome this, we incorporated equivariance properties [21] into the learning objective. The intuition behind this was that the true lesion locations should remain consistent regardless of how we transform the image (e.g., rotation or scaling). When an image was transformed, the corresponding CAM should undergo the same transformation. By enforcing this consistency constraint on CAM through transformation-based regularization, we encouraged the model to focus on actual lesion features rather than arbitrary or position-dependent patterns. This integration promoted robust regularization, ensuring CAM consistency under perturbations (such as affine image transformations), thereby enhancing its spatial alignment with abnormal white matter (WMH) regions.

We implemented the equivalent regularization loss for our volumetric application (see Figure 3). We denoted the backbone network (without global average pooling (GAP) layers)as *θ*(·) and image-transform functions (e.g., scale, rotate, flip, translate) as *π*(·) respectively. For a brain MRI volume X, we had an activation map CAM and visual rating prediction *ŷ* where *CAM* = *θ*(*X*) and *ŷ* = *GAP*(*CAM*). Our equivalent regularization loss could be written as:

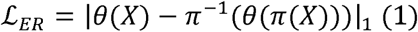

where |… |_1_ denoted the L1 norm. The supervisory signal came from the regression loss for predicting ARWMC visual ratings. We added a global average pooling (GAP) layer at the end of our model to aggregate the CAM of the input image. We adopted mean square error (MSE) loss for both original and transformed images. Our regression loss (MSE loss) could be written as follows:

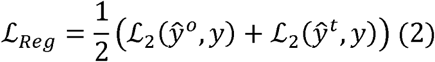

where superscripts o and t represented original and transformed images respectively, and ℒ_2_ denoted the MSE loss. Combining Equitation (1) and (2), our overall training loss was:

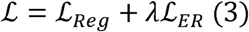

where the λ is the hyper-parameter controled the strength of ℒ*_ER_*.

#### 2.2.2 Extraction of hyperintense map for enhanced WMH boundary delineation

WMH only comprisesed less than 1% of the volume of the whole brain [3] in healthy older subjects and it was dispersed throughout the white matter. Even with enhanced CAM, it could only reflect high-level semantics of the target with a smooth boundary, which made it difficult to delineate a crisp boundary (see Figure 4B). Moreover, we observed that directly using the enhanced CAM as the seed for segmentation, while helpful for large lesions, fell short of capturing the full extent of smaller WMH.

**Figure 4.**
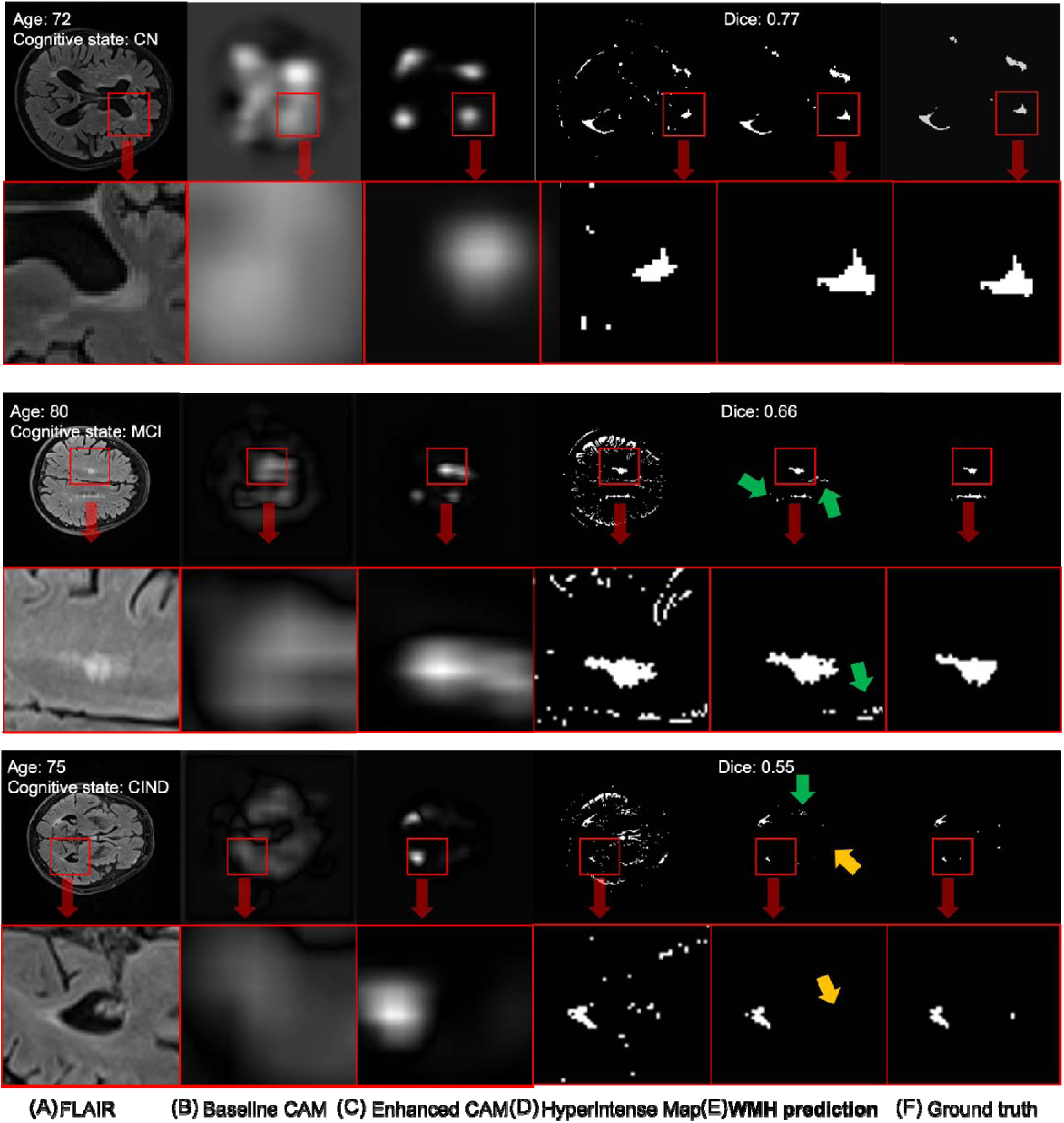
WMH-DualTasker generates high-quality WMH masks by integrating enhanced CAM and Hyperintense Maps. Displayed are three representative cases (with age, disease status and Dice scores shown on each row), each presenting both full axial slices (top) and zoomed-in details (bottom) to enhance visualization. Progressing from left to right, the images showcase (A) FLAIR images, (B) baseline Class Activation Maps (CAM), (C) enhanced CAMs, (D) Hyperintense Maps, (E) WMH predictions from WMH-DualTasker, and finally, (F) the reference standard annotations. The baseline CAM identifies potential WMH regions but with blurry boundaries, in contrast to the enhanced CAM, which delineates these regions with greater precision. Although Hyperintense Maps accurately outline WMH with clear boundaries, they are prone to including false positives. The integration of enhanced CAM with Hyperintense Maps by WMH-DualTasker culminates in the generation of high-quality WMH masks, achieving a similarity to manual annotations and substantially reducing false positives. Green and yellow arrows indicate representative regions of over- and under-segmentation, respectively, when comparing WMH-DualTasker predictions with the ground truth.

To address the challenge posed by the nature of WMH, we extracted a hyperintense map from brain MRI volumes, using the intensity information as prior knowledge for segmentation mask generation (see Equation 4). The hyperintense map (HM) was defined as follows. The values of μ and σ stood for the mean and standard deviation of the grey matter (GM) intensity distribution, which were calculated from a GM mask generated from T2-FLAIR images using FSL-FAST [18]. α was a scaling parameter that adjusts the minimum signal intensity of outliers and i∈Ω denoted all voxel positions in the image space.

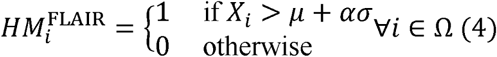

#### 2.2.3 Inference of WMH-DualTasker

To integrate the enhanced CAM and hyperintense map components described above, we first computed the weighted sum of HM and CAM and post-processed with a 3D dense conditional random field (CRF) [22, 23] to generate our final WMH segmentation mask output (see lower half of Figure 3). Importantly, the CRF is a non-learning post-processing technique that refines segmentation boundaries based on spatial and intensity relationships in the input image, with parameters set according to established practices [22, 23]. Unlike trainable components of our model, CRF does not require training or parameter tuning on any dataset.

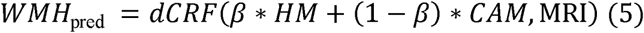

The symbol *dCRF_MRI_* represented the 3D dense CRF function, and the coefficient *β* facilitated the linear combination of HM (Hyperintense Map) and CAM (Class Activation Map), while the MRI image acted as a reference within the CRF algorithm. This integration ensured that the segmentation of WMH was aligned with the specific attributes of each MRI scan. By incorporating the MRI data, the CRF framework harnessed both spatial and intensity information intrinsic to the MRI images. This led to a segmentation that was more contextually enriched.

#### 2.2.4 Model Implementation and Training Details

We used SFCN [20] as our backbone architecture with the last two pooling layers removed for high-resolution CAM. To train the models, we used the AdamW optimizer [24] with an initial learning rate of 0.005 with linear learning rate decay to 0.00005 over 400 epochs. The weight for the equivalent regularization loss, lambda, was initially set to 0. This approach avoided enforcing the equivalent regularization loss at the start of training, which we empirically observed could degrade the quality of the CAM. Starting from the 200th epoch, lambda underwent a stepwise increase, updating every epoch until it reached 0.5. This gradual adjustment supported better training outcomes. We included scale and flip as image transform functions. We set the rescale ratio to 0.5-0.7 and flipped along any axis randomly for each iteration. Standard L2 and L1 loss were used to calculate our regression loss and equivalent regularization loss, respectively. Hyper-parameters alpha and beta were set to be 1.0 and 0.4, respectively. More details about the parameter search range could be found in Table S1 in the supplmentary material.

### 2.3 Evaluation of WMH-DualTasker

We designed our experiments to rigorously evaluate WMH-DualTasker’s performance, understood its components’ contributions, and assessed its potential clinical impact across different populations and cognitive states. Our experimental setup consisted of three main phases: performance evaluation, ablation studies, and downstream analysis (Figure 2).

#### 2.3.1 Performance evaluation and ablation studies

We compared our model’s performance against established supervised methods requiring manual annotation during training: LST-LGA and LST-LPA (Lesion Segmentation Tool’s lesion growth algorithm based on Gaussian mixture modeling and lesion prediction algorithm using logistic regression)[9, 10], UBO-detector (a kNN-based automated pipeline for WMH extraction)[11], Samseg(a Bayesian model-based method for lesion segmentation)[25, 26], 3D full-resolution nnU-Net trained on the MICCAI-WMH dataset (a self-configuring deep learning framework or biomedical image segmentation) [27], and SynthSeg (a CNN-based WMH segmentation approach using only T1-weighted images) [66].

To evaluated the segmentation performance of WMH-DualTasker and other methods (Section 3.2), we adopted five primary metrics: Dice Score and Absolute Volume Difference (AVD), lesion-wise recall, lesion-wise F1, and Intraclass Correlation Coefficient (ICC). The Dice Score measured voxel-wise prediction accuracy for WMH compared to manual annotations, while AVD, a volume-based metric, provided information about a participant’s overall WMH load. The lesion-wise recall and F1 scores evaluated the model’s ability to detect and segment individual WMH lesions defined as 3D connected components, following the criteria established in the MICCAI WMH Segmentation Challenge. The ICC assessed the reliability and agreement between predicted and manual measurements across all subjects. Initial Shapiro-Wilk tests on the MACC-Test dataset revealed that both metrics were not normally distributed (W = 0.89, p < 0.001 for Dice scores; W = 0.87, p < 0.001 for AVD; W = 0.88, p < 0.001 for lesion-wise recall; W = 0.86, p < 0.001 for lesion-wise F1). Therefore, Wilcoxon signed-rank tests with FDR correction were used for comparing methods across all datasets. For visual rating prediction, we rounded the predicted scores to the nearest integer and calculated the mean absolute error (MAE), Pearson correlation coefficient (R), and Kappa score to assess agreement with the the visual rating score from the radiology. To ensure robust evaluation, we reported the experiments with 10 different random seeds and reported the metrics as mean ± standard deviation. Wilcoxon signed-rank tests with FDR correction were employed for comparing ARWMC prediction performance between methods, as the predictions also showed non-normal distribution (W = 0.91, p < 0.001 on MACC-Test dataset).

We conducted ablation studies on the MACC-Test dataset to examine each component’s contribution to WMH-DualTasker (Section 3.3). These studies explored various configurations, including different backbone architectures, CAM methods, and our proposed enhancements such as high-resolution CAM and self-supervised regularization.

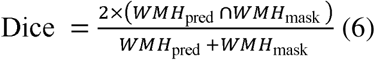

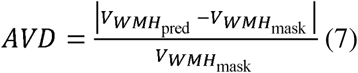

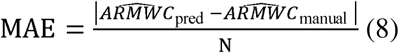

#### 2.3.2 Downstream validation analyses

For downstream validation analyses, we evaluated the utility of WMH volume and ARWMC derived from WMH-DualTasker in predicting cognitive performance across external datasets. We utilized two distinct cohorts: the Alzheimer’s Disease Neuroimaging Initiative (ADNI) database (n=394) as a disease cohort, and the SINGER dataset (n=470) as a community-based cohort. This dual-cohort approach allowed us to investigate the applications of these biomarkers in both clinical and general population settings.

In the ADNI dataset (http://adni.loni.usc.edu), we performed three types of analyses (Figure 5): classification between Cognitively Normal (CN) and Mild Cognitive Impairment (MCI) (155 CN & 146 MCI subjects; Figure 5a), prediction of MCI prognosis (40 MCI non-converters & 40 MCI converters; Figure 5b), and regression analyses of cognitive outcomes (394 subjects; Figure 5c). The regression analyses estimated scores for Mini-Mental State Examination (MMSE), Alzheimer’s Disease Assessment Scale-Cognitive Subscale (ADAS-Cog), and Clinical Dementia Rating Scale sum of boxes (CDRSB). These cognitive measures were selected for their widespread clinical adoption and established relationships with WMH burden in previous studies. MMSE provides a general cognitive assessment that has shown significant correlations with WMH volume [62]. ADAS-Cog, particularly sensitive to attention and processing speed domains, has demonstrated strong associations with WMH burden [63]. CDRSB, which captures both cognitive and functional aspects, has been linked to WMH progression [64, 65].

**Figure 5.**
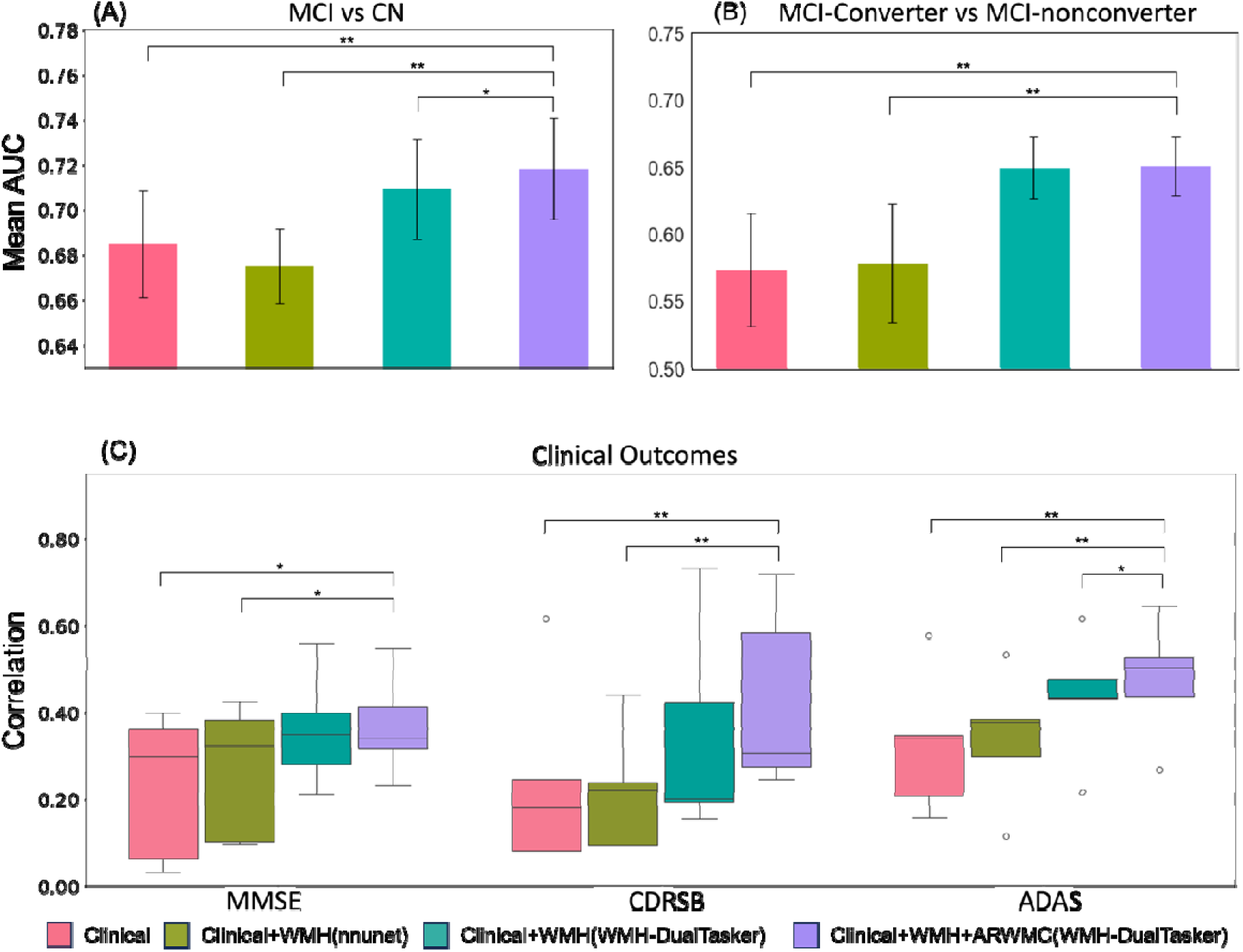
Comparison of prediction models with different combinations of clinical and WMH biomarker features. (Panel A-B) Bar charts showing the mean Area Under the Curve (AUC) values with standard error bars for: (A) classification between Cognitively Normal (CN) and Mild Cognitive Impairment (MCI) subjects and (B) classification of MCI converters versus non-converters. Models compared are clinical factors only (pink), clinical + WMH volume from nnU-Net (olive green), clinical + WMH volume from WMH-DualTasker (teal), and clinical + WMH volume and ARWMC from WMH-DualTasker (purple). (Panel C) Box plots displaying correlation coefficients between predicted and actual cognitive scores for Mini-Mental State Examination (MMSE), Clinical Dementia Rating Scale Sum of Boxes (CDRSB), and Alzheimer’s Disease Assessment Scale-Cognitive Subscale (ADAS-Cog). Each box plot represents the distribution of correlation values obtained from 10-fold cross-validation for each model. Asterisks indicate significant differences between models: *FDR-corrected p < 0.05, **FDR-corrected p < 0.01.

We analyzed these tasks using four different combinations of input variables: clinical and demographic factors only (including age, cardiovascular risk measured by Hachinski Ischemic Scale, Amyloid Standardized Uptake Value Ratio (SUVR), years of education, gender, ApoE4 genotype, and ethnicity), clinical factors plus WMH volume from nnU-Net, clinical factors plus WMH volume from WMH-DualTasker, and clinical factors plus both WMH volume and ARWMC rating from WMH-DualTasker. To ensure robust results, we employed ten-fold nested cross-validation, using logistic regression for classification tasks and kernel ridge regression for cognitive score prediction.

For the SINGER dataset, we explored the relationship between WMH burden and cognitive deterioration, specifically investigating the association between WMH volume predicted by WMH-DualTasker and the (Cardiovascular Risk Factors, Aging, and Incidence of Dementia) CAIDE Dementia Risk Score. The CAIDE score assesses the risk of developing dementia later in life, providing insights into the utility of our biomarkers in evaluating future dementia risk in a general population.

This comprehensive experimental setup allowed us to thoroughly evaluate WMH-DualTasker’s performance, understand its inner workings, and assess its potential clinical impact across various populations and cognitive states.

In the downstream analysis (Section 3.4), the area under the receiver operating characteristic (AUC) curves were calculated to evaluate the predictive capability of different features combinations. We applied ten-fold cross-validation, yielding 10 pairs of AUC values for each comparison between methods. A paired T-test was used to assess statistical significance between methods, with FDR correction applied to account for multiple comparisons. For analysis on the SINGER dataset, where we used Spearman correlation to assess the relationship between WMH volume and CAIDE scores. All statistical analyses were performed using Python version 3.7.11 with the Sklearn package for machine learning metrics and the Statsmodels package for statistical testing.

## 3 Results

Our experiments demonstrated the effectiveness of WMH-DualTasker in WMH segmentation, ARWMC prediction, and clinical applications. Key findings included comparable or superior performance to supervised methods in segmentation tasks, significant improvements in ARWMC prediction, and enhancement in cognitive impairment assessment and dementia risk prediction. We presented detailed results comparing our model to established methods, followed by ablation studies and downstream clinical analyses.

### 3.2 Result for comparative analysis of WMH-DualTasker with segmentation and regression Baseline Models

Table 2 presented the performance of WMH-DualTasker compared to supervised segmentation methods across three datasets. On the MACC-Test dataset, WMH-DualTasker achieved a Dice score of 0.538 ± 0.175, comparable to supervised methods like LST-LPA (0.551 ± 0.230) [10] and superior to LST-LGA (0.403 ± 0.253) [9]. Notably, WMH-DualTasker outperformed all methods in terms of Absolute Volume Difference (AVD), achieving 0.367±0.256 compared to the next best, Samseg [25, 26], at 0.513 ± 0.284.

**Table 2.**
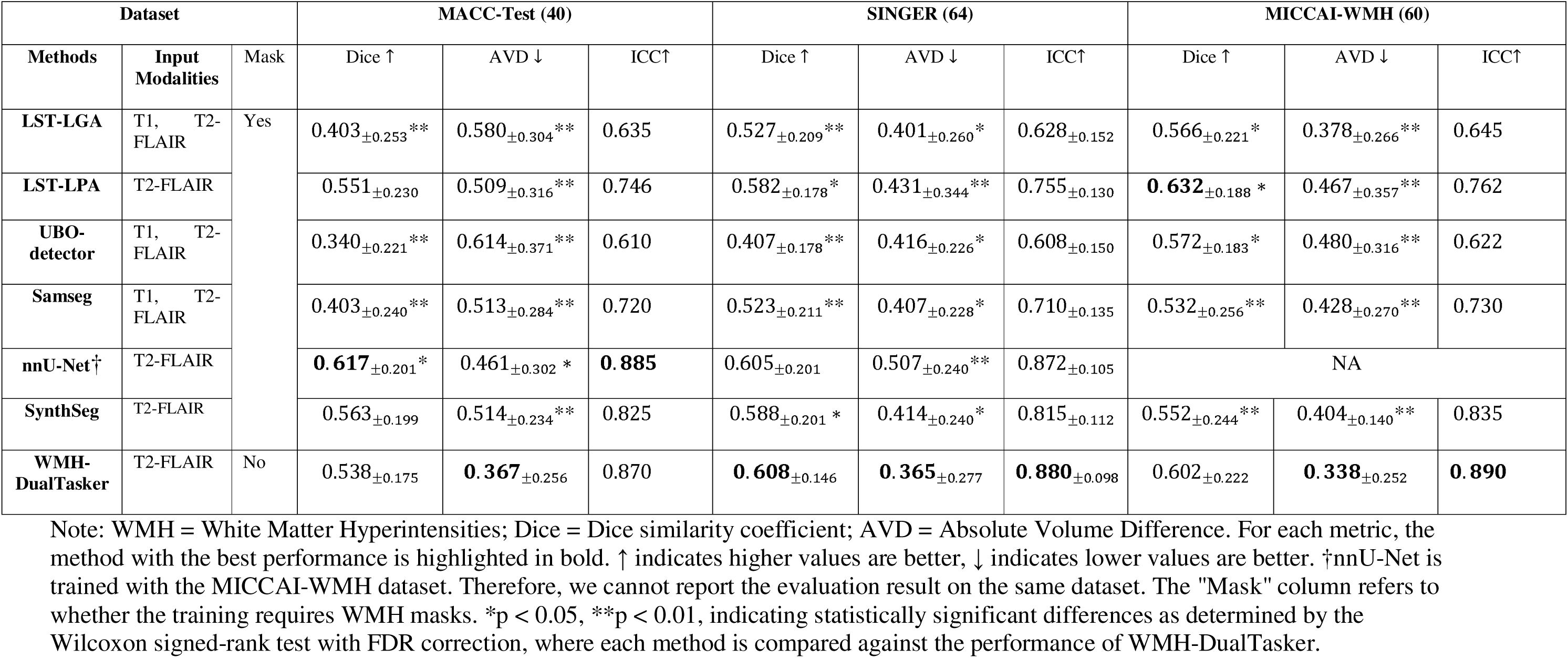
Comparative evaluation of segmentation across all WMH segmentation tools.

On the SINGER dataset, WMH-DualTasker demonstrated the highest Dice score (0.608 ± 0.146) and lowest AVD (0.365 ± 0.277) among all methods, outperforming LST-LPA (Dice: 0.582 ± 0.178, AVD: 0.431 ± 0.344) and UBO-detector (Dice: 0.407 ± 0.178, AVD: 0.416 ± 0.226) [11]. Similarly, for the MICCAI-WMH dataset, our method achieved competitive Dice scores (0.602 ± 0.222) and the best AVD (0.338 ± 0.252), compared to LST-LPA (Dice: 0.632 ± 0.188, AVD: 0.467 ± 0.357).

Although nnU-Net [27] showed a higher Dice score than WMH-DualTasker on the MACC-Test (0.617 ± 0.201 vs 0.538 ± 0.175), WMH-DualTasker achieve better AVD evaluating WMH segmentation [28] (0.367 ± 0.256 vs 0.461 ± 0.302). The superior AVD performance of WMH-DualTasker suggested its better capability in estimating overall WMH burden, which is crucial for clinical applications. The Bland-Altman plots comparing WMH-DualTasker with existing methods across all datasets can be found in Figure S3 in the supplementary material.

Analyzing performance across different brain regions (Tables S7-S9 in the supplementary material), WMH-DualTasker showed consistently strong performance across all anatomical areas. The model performed particularly well in the basal ganglia regions (Dice scores: 0.620-0.635) and frontal lobes (Dice scores: 0.602-0.632), while maintaining robust performance in more challenging areas such as the temporal lobe and infratentorial regions.

Beyond voxel-wise metrics, cluster-level evaluation (Table S10 in the supplementary material) revealed WMH-DualTasker’s superior ability to detect and segment individual lesion clusters. The model achieved higher cluster-wise recall (0.662 ± 0.078 on MACC-Test, 0.632 ± 0.051 on SINGER, and 0.758 ± 0.042 on MICCAI-WMH) and F1 scores (0.582 ± 0.064 on MACC-Test, 0.638 ± 0.050 on SINGER, and 0.677 ± 0.053 on MICCAI-WMH) compared to other methods, demonstrating its effectiveness in identifying both large and small lesion clusters. Visual inspection of the results showed that segmentation (Figure 4E) from WMH-DualTasker can accurately delineate the WMH from healthy tissue and is even very similar to our standard reference (Figure 4F) from our radiologists.

For the ARWMC regression task (Table 3), WMH-DualTasker significantly outperformed standard 3D convolutional backbones [20, 29]. On the MACC-Test dataset, our method achieved a mean absolute error (MAE) of 1.220 ± 1.560, compared to 1.450 ± 1.284 for the next best method SFCN. The Pearson’s correlation (R) of 0.933 ± 0.02 and Cohen’s kappa (κ) of 0.94 ± 0.04 for WMH-DualTasker were substantially higher than all other methods (next best: SFCN with R=0.854±0.03, κ=0.82 ± 0.06), indicating excellent agreement with radiologists’ visual ratings.

**Table 3.**
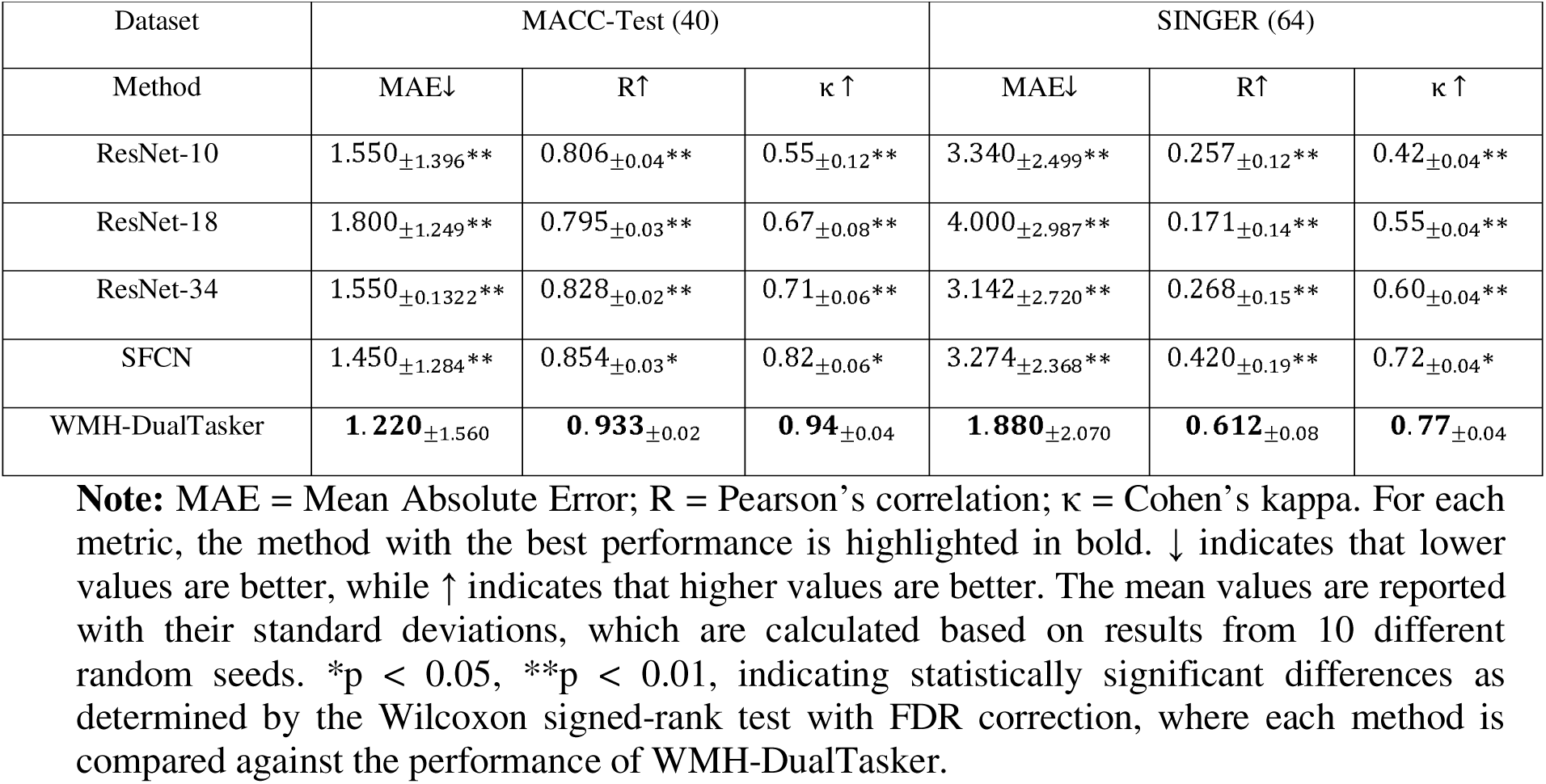
Comparative Evaluation of Regression Across All Methods.

On the SINGER dataset, WMH-DualTasker maintained its superior performance with an MAE of 1.880 ± 2.070, R of 0.612 ± 0.08, and κ of 0.77 ± 0.04, significantly outperforming other methods (next best: SFCN with MAE = 3.274 ± 2.368, R=0.420 ± 0.19, κ=0.72 ± 0.04). These results demonstrated the robust generalization of our method across different datasets and its potential as a reliable tool for automated ARWMC rating.

### 3.3 Result for ablation studies

To assess the effectiveness of each component in our proposed WMH-DualTasker approach, we conducted ablation studies, as shown in Table 4. The results revealed several important insights. First, using only CAM proved insufficient for satisfactory performance (Dice score: 0.133 ± 0.109), underscoring the necessity of the hyperintense map for effective segmentation. Enhancing the resolution of CAM significantly improved segmentation by preserving more details, demonstrating its critical role (Dice score increased to 0.312±0.180). Notably, different backbone networks provided comparable results under this enhancement (e.g., ResNet-10: 0.300 ± 0.152, ResNet-18: 0.283 ± 0.184). Second, the incorporation of self-supervised equivalent regularization loss led to notable improvements in both segmentation and regression performance (Dice: 0.474 ± 0.233, MAE: 1.510 ± 1.660 with scale transformation). This improvement was attributed to the enhanced quality of CAM, which more accurately reflected WMH locations, aiding in more precise ARWMC predictions. Specific affine transformations, such as rotation and flipping, further enhanced performance (Dice: 0.514 ± 0.212, MAE: 1.220 ± 1.560 with scale and flip), although combining all transformations indiscriminately did not yield additional benefits (Dice: 0.466 ± 0.221, MAE: 1.467 ± 1.872 with scale, flip, rotate, and translate). Finally, while CRF provided incremental improvements (final Dice: 0.538±0.175), the use of more advanced CAM methods did not necessarily result in better performance (Grad-CAM: 0.503 ± 0.196, Grad-CAM++: 0.527 ± 0.189). Collectively, these findings highlighted the critical components and configurations that contribute to the efficacy of the WMH-DualTasker model.

**Table 4.**
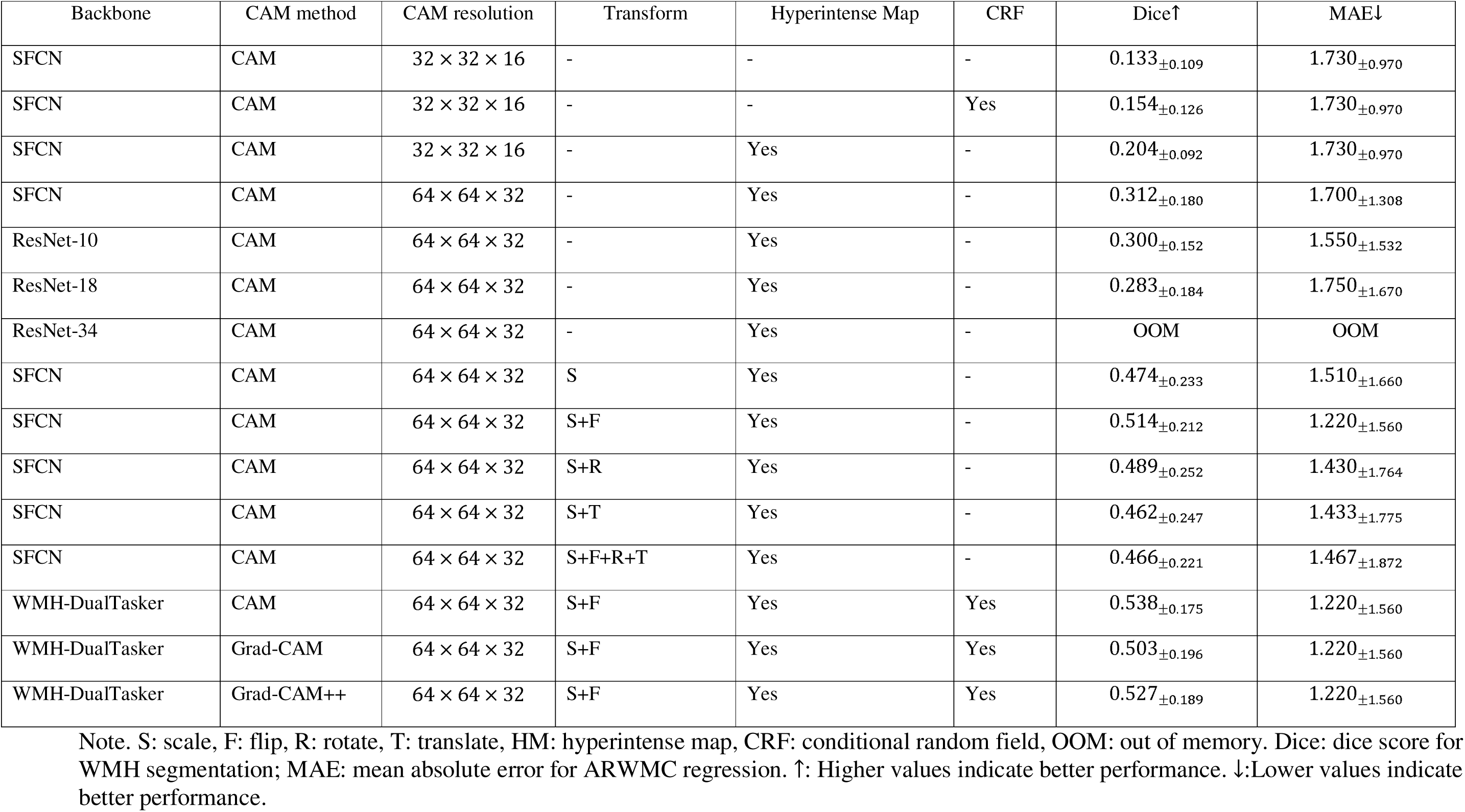
Ablation studies results.

| Backbone | CAM method | CAM resolution | Transform | Hyperintense Map | CRF | Dice↑ | MAE↓ |
| --- | --- | --- | --- | --- | --- | --- | --- |
| SFCN | CAM | $32 \times 32 \times 16$ | - | - | - | $0.133_{\pm 0.109}$ | $1.730_{\pm 0.970}$ |
| SFCN | CAM | $32 \times 32 \times 16$ | - | - | Yes | $0.154_{\pm 0.126}$ | $1.730_{\pm 0.970}$ |
| SFCN | CAM | $32 \times 32 \times 16$ | - | Yes | - | $0.204_{\pm 0.092}$ | $1.730_{\pm 0.970}$ |
| SFCN | CAM | $64 \times 64 \times 32$ | - | Yes | - | $0.312_{\pm 0.180}$ | $1.700_{\pm 1.308}$ |
| ResNet-10 | CAM | $64 \times 64 \times 32$ | - | Yes | - | $0.300_{\pm 0.152}$ | $1.550_{\pm 1.532}$ |
| ResNet-18 | CAM | $64 \times 64 \times 32$ | - | Yes | - | $0.283_{\pm 0.184}$ | $1.750_{\pm 1.670}$ |
| ResNet-34 | CAM | $64 \times 64 \times 32$ | - | Yes | - | OOM | OOM |
| SFCN | CAM | $64 \times 64 \times 32$ | S | Yes | - | $0.474_{\pm 0.233}$ | $1.510_{\pm 1.660}$ |
| SFCN | CAM | $64 \times 64 \times 32$ | S+F | Yes | - | $0.514_{\pm 0.212}$ | $1.220_{\pm 1.560}$ |
| SFCN | CAM | $64 \times 64 \times 32$ | S+R | Yes | - | $0.489_{\pm 0.252}$ | $1.430_{\pm 1.764}$ |
| SFCN | CAM | $64 \times 64 \times 32$ | S+T | Yes | - | $0.462_{\pm 0.247}$ | $1.433_{\pm 1.775}$ |
| SFCN | CAM | $64 \times 64 \times 32$ | S+F+R+T | Yes | - | $0.466_{\pm 0.221}$ | $1.467_{\pm 1.872}$ |
| WMH-DualTasker | CAM | $64 \times 64 \times 32$ | S+F | Yes | Yes | $0.538_{\pm 0.175}$ | $1.220_{\pm 1.560}$ |
| WMH-DualTasker | Grad-CAM | $64 \times 64 \times 32$ | S+F | Yes | Yes | $0.503_{\pm 0.196}$ | $1.220_{\pm 1.560}$ |
| WMH-DualTasker | Grad-CAM++ | $64 \times 64 \times 32$ | S+F | Yes | Yes | $0.527_{\pm 0.189}$ | $1.220_{\pm 1.560}$ |

Note. S: scale, F: flip, R: rotate, T: translate, HM: hyperintense map, CRF: conditional random field, OOM: out of memory. Dice: dice score for WMH segmentation; MAE: mean absolute error for ARWMC regression. ↑: Higher values indicate better performance. ↓: Lower values indicate better performance.

These findings highlighted the critical components and configurations that contribute to the efficacy of the WMH-DualTasker model, with the combination of high-resolution CAM, hyperintense map, and self-supervised regularization providing the optimal performance.

### 3.4 Result for downstream validation analyses

#### 3.4.1 Downstream analysis in AD cohort (ADNI dataset)

Our analysis revealed that incorporating WMH volume significantly enhanced the accuracy of cognitive impairment classification, prediction of disease progression, and estimation of cognitive performance scores (Figure 5).

The addition of WMH volume derived from WMH-DualTasker to clinical variables substantially improved the discrimination between Mild Cognitive Impairment (MCI) and Cognitively Normal (CN) subjects, increasing the Area Under the Curve (AUC) from 0.681 to 0.711 (p < 0.001). Combining both WMH volume and ARWMC scores further improved classification performance, achieving the highest AUC of 0.718 (Figure 5A).

WMH-DualTasker-derived biomarkers also enhanced the prediction of MCI conversion, with the AUC increasing from 0.618 to 0.652 (p < 0.001) for distinguishing between MCI converters and non-converters. This improvement underscored the potential of WMH features in identifying individuals at higher risk of progressing to dementia (Figure 5B).

Importantly, WMH volume predictions from WMH-DualTasker consistently outperformed those from nnUNet across both classification tasks, highlighting the robustness of our weakly supervised approach.

For continuous clinical outcomes (MMSE, CDRSB, and ADAS scores), incorporating WMH biomarkers consistently improved the prediction performance as measured by correlation coefficients between the predicted scores from kernel ridge regression and the actual clinical scores. The most notable improvements were observed for CDRSB and ADAS predictions, where the combination of clinical variables with WMH volume and ARWMC scores from WMH-DualTasker yielded the highest correlations (Figure 5C).

#### 3.4.2 Downstream analysis in a community-based cohort (SINGER dataset)

To assess the utility of WMH-DualTasker in a general population setting, we analyzed the SINGER dataset, revealing a significant positive correlation between WMH volume predicted by WMH-DualTasker and the CAIDE Dementia Risk Score [30–32] (Spearman’s rho = 0.19, p < 0.001, Figure 6). This finding supported the potential utility of WMH volume as a biomarker for future dementia risk in community-based populations.

**Figure 6.**
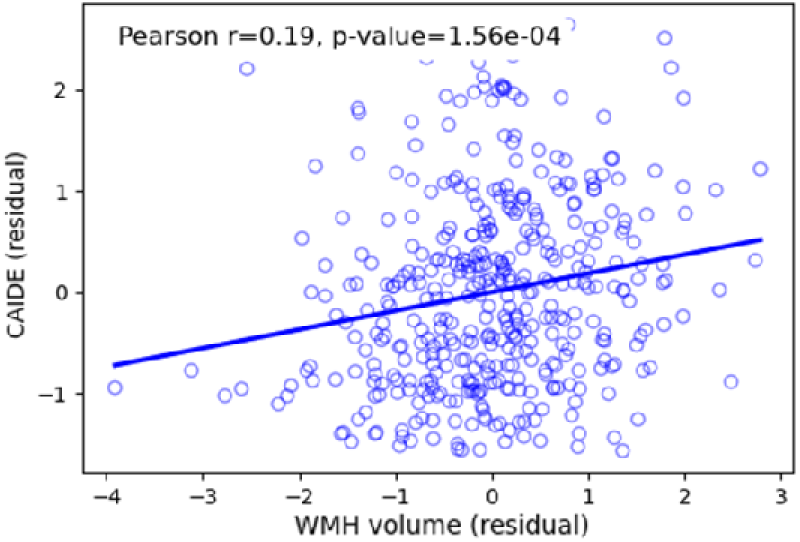
Higher WMH volumes are associated with increased dementia risk in a community-based cohort. Scatter plot illustrating the positive correlation between WMH volume predicted by WMH-DualTasker and CAIDE Risk Score in the SINGER cohort. This relationship suggests that increased WMH burden, as quantified by our model, is linked to higher risk of future dementia in a general population setting. CAIDE: Cardiovascular Risk Factors, Aging, and Incidence of Dementia.

The observed correlation, while modest, was clinically meaningful given the complex, multifactorial nature of dementia risk. It suggested that WMH-DualTasker can provide valuable information for assessing cognitive health in general populations, complementing traditional risk factors captured by the CAIDE score [33]. These findings, combined with the results from the ADNI cohort, demonstrated the versatility and clinical relevance of WMH-DualTasker across different population types and cognitive states.

### 3.5 Usage of WMH-DualTasker

The usage of WMH-DualTasker was relatively straightforward. Our GitHub repository provided examples to guide users for applying WMH-DualTasker. Our wrapper function will automatically perform the pre-processing for inputs with arbitrary resolution; training requires around 8GB of GPU memory on an RTX3090. Inference on a single subject is computationally efficient, requiring only a forward pass through the Deep Neural Network (DNN) followed by the application of Conditional Random Fields (CRF). This process is extremely fast, typically completed in a matter of seconds on standard GPU hardware and a few minutes on CPU.

## 4. Discussion

In this study, we present WMH-DualTasker, a deep learning-based algorithm that simultaneously performs white matter hyperintensity (WMH) segmentation and visual rating prediction. Our main contribution is the development of a model that requires only visual rating scales for training, eliminating the need for labor-intensive, voxel-wise WMH annotations. This approach significantly reduces the time and expertise required for data preparation, making it particularly suitable for large-scale brain imaging studies. WMH-DualTasker employs a technique that leverages activation maps from the regression task to guide WMH segmentation. We enhance this process by incorporating self-supervised regularization and integrating a hyperintense map, improving lesion localization and sensitivity. By balancing accuracy, efficiency, and clinical relevance, our approach advances automated WMH analysis, offering a comprehensive tool for assessing cerebrovascular health in both research and clinical settings. The dual functionality of WMH-DualTasker in providing both segmentation and visual ratings makes it a versatile solution for quantifying WMH burden, potentially accelerating research in neurodegenerative and cerebrovascular diseases.

Recent research has increasingly emphasized the significance of WMH as a key marker of cerebrovascular health, with studies like FINGER demonstrating its value in predicting cognitive outcomes [68]. The quantification of WMH volume has emerged as a critical metric in large-scale clinical trials and population studies [69, 70], highlighting the need for efficient assessment tools. However, existing WMH segmentation methods have limitations. Unsupervised methods [34–38] leverage the inherent MRI data characteristics, but often lack precision in identifying subtle WMH nuances[59, 60], particularly in cases with low lesion load or early-stage disease [57]. Traditional supervised deep learning approaches [9, 11–13, 24, 39–42] have shown promising results in recent studies [12, 13], however they rely on manually annotated WMH masks, which are time-consuming and expensive to produce, typically requiring 30-120 minutes per scan by trained experts [13]. This dependency on extensive manual annotation makes these methods impractical for large-scale applications. Recent developments in semi-supervised learning have shown potential to reduce annotation burden while maintaining performance [67]. Alternative weakly supervised methods have explored various forms of annotations, such as bounding boxes [45, 46] and partial annotations [47–53]. While these approaches have shown promise [54], they face particular challenges with WMH segmentation due to the dispersed nature of the lesions, as WMH often appears as multiple, scattered abnormalities throughout the white matter [3, 4], with lesion sizes varying from millimeters to centimeters [4]. Moreover, bounding boxes and scribbles are ill-suited for capturing this scattered distribution without extensive manual effort.

To address these limitations, WMH-DualTasker leveraged the Age-Related White Matter Changes (ARWMC) visual rating scale only as a supervisory signal. By utilizing activation maps generated from the regression task as initialization for WMH segmentation, WMH-DualTasker can significantly reduce the time and expertise required for data preparation. To enhance the localization capability of the activation map, we introduce an equivalent regularization loss which ensures that it learns transformation invariance, increasing its sensitivity to less distinguishable lesion regions while suppressing activations in non-lesion areas.

We evaluated WMH-DualTasker on both internal and external datasets. Our results demonstrated that WMH-DualTasker achieves performance comparable or superior to the existing supervised methods in WMH segmentation on both internal (MACC) and external (MICCAI-WMH and SINGER) validations. This is particularly noteworthy given that WMH-DualTasker operates with only image-level labels, whereas the existing supervised methods require pixel-level annotations. Notably, while fully supervised methods like nn-UNet achieve higher Dice scores in some cases, WMH-DualTasker demonstrates superior performance in Absolute Volume Difference (AVD), a metric more directly leveraged in research analysis. Moreover, WMH-DualTasker can jointly predict visual rating scores. We observed good to high Kappa agreement concerning the radiologists’ ratings on the external validation (SINGER). The impact of training data size on model performance is illustrated in Figure S2, showing that at least 400 training samples are needed for optimal performance. This indicates WMH-DualTasker’s potential to serve as a valuable reference tool for clinical experts. Overall, our extensive evaluation across multiple datasets confirms that WMH-DualTasker consistently produces accurate WMH volume measurements and ARWMC scores, establishing it as a reliable and generalizable tool for assessing WMH across both Asian and Caucasian populations.

Moreover, we demonstrated the clinical utility of WMH-DualTasker’s outputs in both AD-specific (ADNI) and community-based (SINGER) settings. The incorporation of WMH biomarkers derived from WMH-DualTasker significantly enhanced the assessment of cognitive impairment and prediction of disease progression. Specifically, combining both WMH volume and ARWMC scores improved the accuracy of MCI/CN classification and MCI conversion prediction. The consistent improvements across various cognitive measures and prediction tasks underscore our approach’s clinical value. In the community-cohort (SINGER), we found a correlation between WMH volume and the CAIDE Dementia Risk Score, indicating the relevance of WMH-DualTasker’s outputs in assessing future dementia risk in a general population. These findings collectively demonstrate the potential of WMH-DualTasker as a valuable tool for more accurate cognitive assessment and prognosis in both research and clinical settings.

Given these compelling results, WMH-DualTasker shows great potential for integration into radiology and research workflows. Users can directly apply the pre-trained model (from our github repo) to their T2-FLAIR MRI images without fine-tuning, obtaining rapid and consistent WMH segmentations and ARWMC scores. This out-of-the-box functionality is particularly valuable for clinical settings or large-scale studies where manual annotations are impractical. For instance, a hospital could implement WMH-DualTasker to assist radiologists in assessing stroke risk or monitoring cerebrovascular disease progression across numerous patients. Researchers conducting population-wide studies could efficiently process thousands of MRI scans to investigate relationships between WMH burden and various neurological conditions or cognitive decline. If users wish to optimize performance for their specific dataset, they can fine-tune the model using only visual rating scores for a small subset of their data, without the need for time-consuming manual WMH annotations. Our GitHub repository provides comprehensive documentation and scripts for both direct application and fine-tuning, ensuring WMH-DualTasker can be easily integrated into diverse clinical and research environments.

Our study has several limitations. While we utilized the ARWMC scale due to its availability in our datasets, we did not validate other clinically prevalent scales, such as the Fazekas scale [7]. This analysis focused on cross-sectional data, and future research should explore the algorithm’s performance on longitudinal data to assess WMH progression over time. Additionally, further clinical validation through prospective studies in real-world settings is essential to evaluate the model’s performance in routine clinical workflows and its impact on clinical decision-making. Moving forward, our model architecture could potentially be adapted to different visual rating scales in future studies. While we currently use ARWMC (range 0-30), adaptation to the Fazekas scale (range 0-3) would require consideration of the model’s discriminative capabilities at a more condensed scale range. The adaptation would primarily involve modifying the regression head of our network to predict Fazekas scores while maintaining the same segmentation architecture. However, the effectiveness of such adaptation may vary depending on the granularity of the rating scale - more condensed scoring ranges could present challenges for the model’s discriminative ability. We call for further validation of our approach across different visual rating scales and clinical settings. These validation steps will help position WMH-DualTasker as a reliable tool for assessing and managing cerebrovascular disorders associated with WMH, offering valuable assistance to healthcare professionals.

## 5. Conclusion

In conclusion, WMH-DualTasker, a novel dual-task deep learning model with self-supervised consistency, achieved accurate and generalizable performance for automatic White Matter Hyperintensity (WMH) segmentation and visual rating prediction using T2-FLAIR MRI. This weakly-supervised approach significantly reduces the annotation burden by utilizing only image-level visual ratings for training, while effectively performing both voxel-wise segmentation and global ARWMC score prediction. The model’s robustness was demonstrated across multiple datasets, including both disease-specific and community-based cohorts. Considering its effectiveness in joint segmentation and regression tasks, and its proven clinical utility in cognitive impairment assessment and dementia risk prediction, WMH-DualTasker represents a valuable tool for both clinical practice and scientific research. Its ability to process large-scale datasets efficiently makes it particularly suited for population-wide studies, potentially advancing our understanding of cerebral small vessel disease and cognitive disorders.

## Acknowledgements

We extend our heartfelt gratitude to all the participants and the dedicated research teams at MACC and SINGER for their invaluable contributions. A special thanks goes to Dr. Hugo Kuijf for not only hosting the MICCAI-WMH segmentation challenge but also for making the dataset widely accessible to the public. Additionally, we are deeply appreciative of the Alzheimer’s Disease Neuroimaging Initiative (ADNI) for generously making their dataset available to the research community. These collaborative efforts are fundamental to the advancement of our work.

Data collection and sharing for this project was funded by the Alzheimer’s Disease Neuroimaging Initiative (ADNI) (National Institutes of Health Grant U01 AG024904) and DOD ADNI (Department of Defense award number W81XWH-12-2-0012). ADNI is funded by the National Institute on Aging, the National Institute of Biomedical Imaging and Bioengineering, and through generous contributions from the following: AbbVie, Alzheimer’s Association; Alzheimer’s Drug Discovery Foundation; Araclon Biotech; BioClinica, Inc.; Biogen; Bristol-Myers Squibb Company; CereSpir, Inc.; Cogstate; Eisai Inc.; Elan Pharmaceuticals, Inc.; Eli Lilly and Company; EuroImmun; F. Hoffmann-La Roche Ltd and its affiliated company Genentech, Inc.; Fujirebio; GE Healthcare; IXICO Ltd.; Janssen Alzheimer Immunotherapy Research & Development, LLC.; Johnson & Johnson Pharmaceutical Research & Development LLC.; Lumosity; Lundbeck; Merck & Co., Inc.; Meso Scale Diagnostics, LLC.; NeuroRx Research; Neurotrack Technologies; Novartis Pharmaceuticals Corporation; Pfizer Inc.; Piramal Imaging; Servier; Takeda Pharmaceutical Company; and Transition Therapeutics. The Canadian Institutes of Health Research is providing funds to support ADNI clinical sites in Canada. Private sector contributions are facilitated by the Foundation for the National Institutes of Health (www.fnih.org). The grantee organization is the Northern California Institute for Research and Education, and the study is coordinated by the Alzheimer’s Therapeutic Research Institute at the University of Southern California. ADNI data are disseminated by the Laboratory for Neuro Imaging at the University of Southern California.

## Declarations

decision-making related to the acceptance of this article for publication.

## Ethics approval and consent

The local Institutional Review Boards at each participating institution approved participant recruitment and data collection for the ADNI project. Informed written consent from participants was obtained at each participating site. For up-to-date information, see www.adni-info.org.

## Consent for publication

All co-authors consent to publish the manuscript.

