## supplemental material for "WMH-DualTasker: A weakly-supervised deep learning model for automated white matter hyperintensities segmentation and visual rating prediction"

**Appendix overview**

**Supplementary Figure S1 Network configuration …………………………………………………………….…..2**

**Supplementary Figure S2 Impact of the size of training data on WMH-DualTasker performance ……….…2**

**Supplementary Figure S3 Bland-Altman Plots for WMH-DualTasker vs Baseline Comparisons………...….…3**

**Supplementary Table S1 Hyperparameters search ranges for WMH-DualTasker on the validation set ……4**

**Supplementary Table S2 Demographic information of Memory, Ageing and Cognition Centre (MACC) dataset ………………………………………………………………………………………………………...…..….5**

**Supplementary Table S3 Demographic information of subjects from SINgapore GERiatric (SINGER) Intervention Study ………………………………………………………...……………………………………..….5**

**Supplementary Table S4 Demographic information of CN and MCI subjects from Alzheimer's Disease Neuroimaging Initiative (ADNI) datasets …………………………………….………………………………..….6**

**Supplementary Table S5 Demographic information of MCI prognosis subjects from Alzheimer's Disease Neuroimaging Initiative (ADNI) datasets ………………………………………………………………….…..….6**

**Supplementary Table S6 Ablation studies on alpha and beta for conditional random field (CRF) ..………… 7**

**Supplementary Table S7 Region-level evaluation across all WMH segmentation tool (MACC) .…………….. 8**

**Supplementary Table S8 Region-level evaluation across all WMH segmentation tool (SINGER) .…………... 9**

**Supplementary Table S9 Region-level evaluation across all WMH segmentation tool (MICCAI-WMH) …...10**

**Supplementary Table S10 Cluster-level evaluation across all WMH segmentation tool …………..…………..11**

**Figure S1. Network configuration.** The underlying deep learning model of the WMH-DualTasker is relatively straightforward. It consists of three main components arranged in sequence. The first component, repeated three times (x3), contains four sequential layers: Conv3D, BatchNorm, MaxPool2x2x2, and ReLU. The second component, repeated twice (x2), has a similar structure but omits the MaxPool layer, which consists of Conv3D, BatchNorm, and ReLU. The third component contains just two layers: AvgPool and Linear. Regarding inference speed, the model takes approximately 2 seconds on a GTX3090 GPU and less than 1 minute on a CPU.

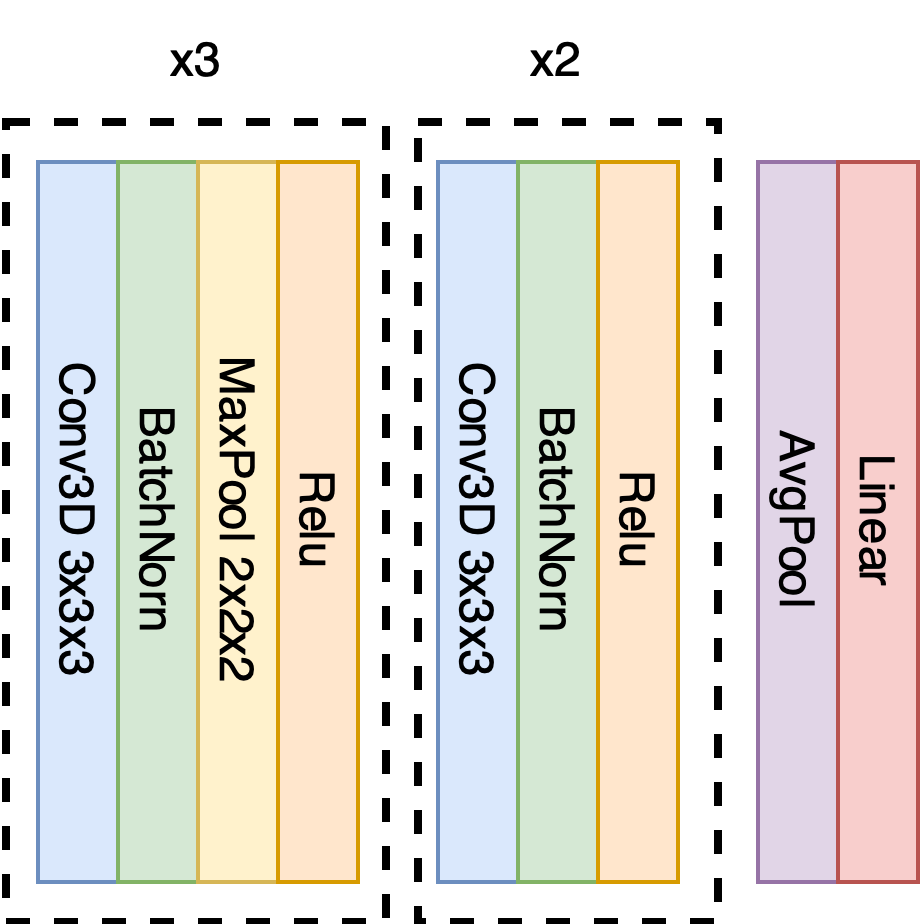

**Figure S.2 Impact of the size of training data on WMH-DualTasker performance.** On the left panel, segmentation accuracy improves as demonstrated by ascending Dice scores. On the right, regression precision is shown through decreasing Mean Absolute Error (MAE) values. The x-axis, representing the number of training samples, indicates that more training data directly enhances WMH-DualTasker's accuracy in both segmentation and regression.

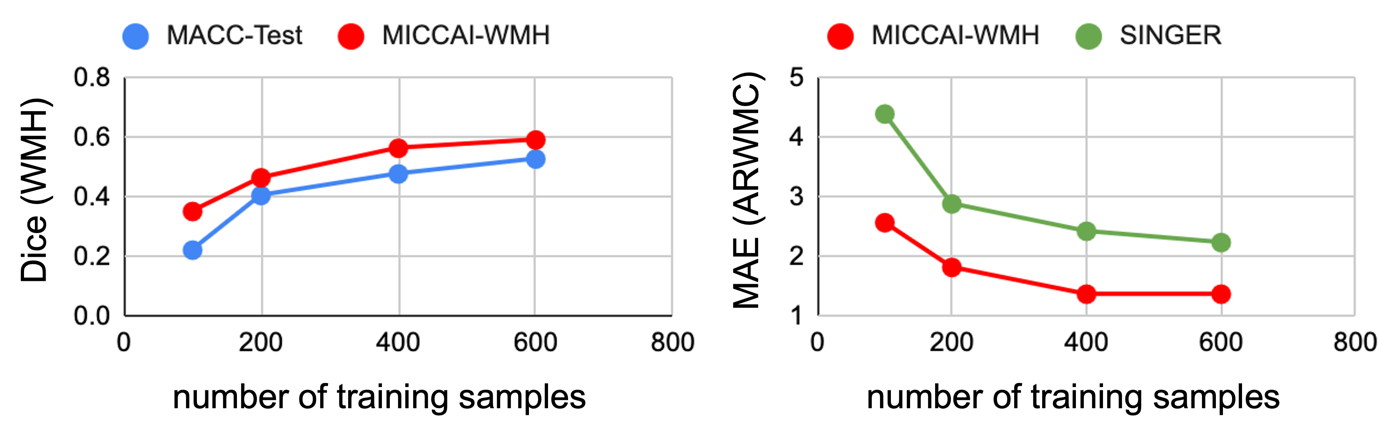

An important consideration is the number of subjects (see Figure B) required for effective training. Our evaluation indicates that WMH-DualTasker requires at least 400 training samples to achieve optimal performance, with smaller datasets significantly reducing the accuracy of both segmentation and regression tasks.

**Figure S.3. Bland-Altman plots comparing WMH-DualTasker with existing segmentation methods** (LST-LGA, LST-LPA, UBO-detector, Samseg, and nnU-Net) across SINGER, MACC-Test, and MICCAI-WMH datasets. Each plot shows volume differences (method prediction minus ground truth) against ground truth volumes (log scale). Green dots represent WMH-DualTasker predictions, and red dots represent baseline method predictions. Horizontal lines indicate mean difference (solid) and 95% limits of agreement (dashed) for each method. Lower spread of differences indicates better agreement with ground truth.
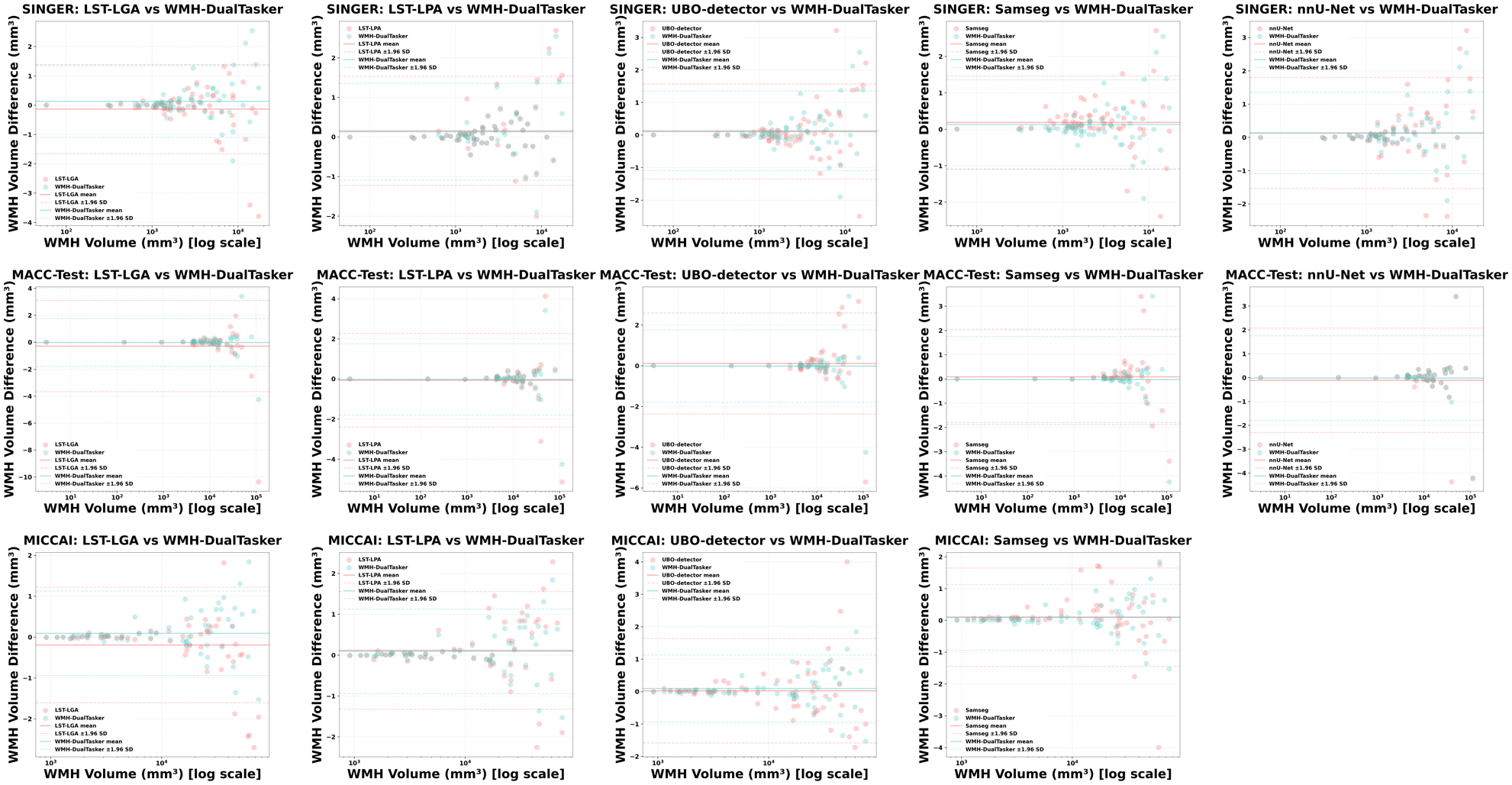

**Table S.1 Hyperparameters search ranges for WMH-DualTasker on the validation set.** To optimize the hyperparameters of the WMH-DualTasker model on the validation set, we employed Optuna (<https://github.com/optuna/optuna>), an open-source framework specifically designed for hyperparameter optimization. In the table, the "Search Range" column outlines the range of values that were explored during the optimization process for each hyperparameter, setting the minimum and maximum boundaries within which Optuna searched for the optimal value. The "Specification" column presents the final values chosen by Optuna after assessing the model's performance on the validation set.

| **Hyperparameter** | **Search Range** | **Specification** |
| --- | --- | --- |
| Learning rate | 0.00001 – 0.1 | 0.005 |
| Learning rate decay | 0.00005 – 0.01 | 0.00005 |
| Learning rate steps | 100 – 1000 | 200 |
| Batch Size | 16 – 128 | 64 |
| Epochs | 100 – 1000 | 400 |
| $\lambda$ | 0.01 – 10 | 1.0 |
| $\beta$ | 0.01 – 1 | 0.4 |

**Table S2 Demographic information of Memory, Ageing and Cognition Centre (MACC) dataset.**

|  | **Training Split**  **(n = 600)** | **Validation Split**  **(n = 40)** | **Test split**  **(n = 20)** | ***p*** |
| --- | --- | --- | --- | --- |
| Age, mean (SD), years | 73.4 (5.2) | 74.1(4.9) | 73.6 (4.7) | 0.68 |
| Gender, Female/Male | 296/304 | 21/19 | 9/11 | 0.32 |
| Ethnicity, Chinese/Non | 383/17 | 34/6 | 17/3 | 0.77 |
| Education, mean (SD), years | 8.3 (4.8) | 8.2 (4.4) | 8.1 (4.5) | 0.53 |
| CDR-SOB, mean (SD) | 0.53 (0.11) | 0.47 (0.09) | 0.50 (0.07) | 0.84 |
| Cardiovascular Risk, means (SD) | 1.7 (1.0) | 1.6 (1.0) | 1.6 (1.0) | 0.80 |
| MMSE, mean (SD) | 28.9 (1.1) | 29.1 (1.0) | 29.1 (1.0) | 0.83 |
| ADAS-Cog score, mean (SD) | 9.7 (5.1) | 9.8 (4.8) | 9.5 (4.2) | 0.76 |
| ARWMC, mean (SD) | 6.7 (2.2) | 6.6 (1.9) | 6.4 (1.7) | 0.59 |

**Table S3. Demographic information of subjects from SINgapore GERiatric (SINGER) Intervention Study.**

|  | **SINGER**  **(n = 64)** | **SINGER***  **(n = 470)** |
| --- | --- | --- |
| Age, mean (SD), years | 74.3 (4.4) | 74.2 (4.9) |
| Gender, Female/Male | 34/30 | 249/221 |
| Ethnicity, Chinese/Non | 62/2 | 458/12 |
| Education, mean (SD), years | 9.1(3.3) | 9.0(3.4) |
| CDR-SOB, mean (SD) | 0.41 (0.2) | 0.37 (0.1) |
| Cardiovascular Risk, means (SD) | 1.6 (1.1) | 1.5 (0.8) |
| CAIDE, mean (SD) | 1.22 (0.13) | 1.19 (0.12) |
| ARWMC, mean (SD) | 5.1 (1.3) | 4.8 (0.9) |

**Table S4. Demographic information of CN and MCI subjects from Alzheimer's Disease Neuroimaging Initiative (ADNI) datasets.**

|  | **CN**  **(n = 155)** | **MCI**  **(n = 146)** | ***p*** |
| --- | --- | --- | --- |
| Age, mean (SD), years | 73.1 (7.5) | 71.9 (7.9) | 0.13 |
| Gender Female/Male | 80/75 | 76/70 | 0.09 |
| Ethnicity, Chinese/Non | 5/150 | 3/143 | 0.15 |
| Education, mean (SD), years | 16.0(2.6) | 17.1 (2.4) | 0.20 |
| Cardiovascular Risk (SD) | 1.5 (1.0) | 1.7 (1.1) | **0.03** |
| MMSE score, mean (SD) | 29.1 (1.4) | 28.3 (2.0) | **<0.01** |
| ADAS-Cog score, mean (SD) | 6.0 (3.2) | 10.1 (4.2) | **<0.01** |
| CDR sum, mean (SD) | 0.12 (0.43) | 1.4 (1.0) | **<0.01** |
| WMHs, cm3 | 8.22 (13.98) | 9.34 (12.77) | **0.02** |

**Table S5. Demographic information of MCI prognosis** **subjects from Alzheimer's Disease Neuroimaging Initiative (ADNI) datasets.**

|  | **MCI-converter**  **(n = 40)** | **MCI-nonconverter**  **(n = 40)** | ***p*** |
| --- | --- | --- | --- |
| Age, mean (SD), years | 71.8 (7.7) | 71.9 (7.5) | 0.45 |
| Gender Female/Male | 22/18 | 23/17 | 0.39 |
| Ethnicity, Chinese/Non | 2/38 | 0/40 | 0.56 |
| Education, mean (SD), years | 17.0 (2.0) | 17.2 (2.8) | 0.82 |
| Cardiovascular Risk (SD) | 1.6 (0.9) | 1.7 (1.1) | 0.49 |
| MMSE score, mean (SD) | 28.3 (1.5) | 28.1 (1.9) | 0.39 |
| ADAS-Cog score, mean (SD) | 10 (4.2) | 10 (4.2) | 0.21 |
| CDR sum, mean (SD) | 1.5 (1.0) | 1.5 (1.1) | 0.21 |
| WMHs, cm3 | 9.25 (10.68) | 9.65 (13.31) | 0.19 |

**Table S6. Ablation studies on alpha and beta for conditional random field (CRF) on MACC-Test.**

| $\alpha$ | $\beta$ | Dice$\uparrow$ | AVD $\downarrow$ | ICC$\uparrow$ |
| --- | --- | --- | --- | --- |
| 0.6 | 0.2 | ${0.485}_{\pm0.195}$ | ${0.420}_{\pm0.310}$ | $0.795$ |
| 0.8 | 0.2 | ${0.510}_{\pm0.182}$ | ${0.405}_{\pm0.290}$ | $0.812$ |
| 1.0 | 0.2 | ${0.528}_{\pm0.175}$ | ${0.395}_{\pm0.280}$ | $0.825$ |
| 1.2 | 0.2 | ${0.520}_{\pm0.187}$ | ${0.397}_{\pm0.226}$ | $0.844$ |
| 1.5 | 0.2 | $\mathbf{0.541}_{\boldsymbol{\pm0.155}}$ | ${0.400}_{\pm0.295}$ | $0.818$ |
| 0.6 | 0.3 | ${0.515}_{\pm0.192}$ | ${0.415}_{\pm0.305}$ | $0.810$ |
| 0.8 | 0.3 | ${0.515}_{\pm0.190}$ | ${0.410}_{\pm0.300}$ | $0.805$ |
| 1.0 | 0.3 | ${0.540}_{\pm0.178}$ | ${0.380}_{\pm0.265}$ | $\mathbf{0.845}$ |
| 1.2 | 0.3 | ${0.530}_{\pm0.185}$ | ${0.390}_{\pm0.280}$ | $0.835$ |
| 1.5 | 0.3 | ${0.525}_{\pm0.195}$ | ${0.405}_{\pm0.300}$ | $0.825$ |
| 0.6 | 0.4 | ${0.495}_{\pm0.202}$ | ${0.440}_{\pm0.320}$ | $0.770$ |
| 0.8 | 0.4 | ${0.525}_{\pm0.185}$ | ${0.395}_{\pm0.285}$ | $0.825$ |
| **1.0** | **0.4** | ${0.538}_{\pm0.175}$ | $\mathbf{0.367}_{\pm0.256}$ | $0.844$ |
| 1.2 | 0.4 | ${0.535}_{\pm0.182}$ | ${0.375}_{\pm0.270}$ | $0.840$ |
| 1.5 | 0.4 | ${0.530}_{\pm0.192}$ | ${0.390}_{\pm0.280}$ | $0.832$ |
| 0.6 | 0.5 | ${0.470}_{\pm0.215}$ | ${0.450}_{\pm0.330}$ | $0.760$ |
| 0.8 | 0.5 | ${0.505}_{\pm0.190}$ | ${0.420}_{\pm0.300}$ | $0.795$ |
| 1.0 | 0.5 | ${0.530}_{\pm0.180}$ | ${0.385}_{\pm0.270}$ | $0.830$ |
| 1.2 | 0.5 | ${0.520}_{\pm0.185}$ | ${0.400}_{\pm0.285}$ | $0.820$ |
| 1.5 | 0.5 | ${0.510}_{\pm0.200}$ | ${0.415}_{\pm0.295}$ | $0.810$ |
| 0.6 | 0.6 | ${0.460}_{\pm0.220}$ | ${0.445}_{\pm0.340}$ | $0.750$ |
| 0.8 | 0.6 | ${0.490}_{\pm0.205}$ | ${0.430}_{\pm0.310}$ | $0.785$ |
| 1.0 | 0.6 | ${0.520}_{\pm0.190}$ | ${0.400}_{\pm0.285}$ | $0.820$ |
| 1.2 | 0.6 | ${0.510}_{\pm0.195}$ | ${0.410}_{\pm0.295}$ | $0.810$ |
| 1.5 | 0.6 | ${0.505}_{\pm0.210}$ | ${0.425}_{\pm0.310}$ | $0.805$ |

Table S7 **Region-level evaluation across all WMH segmentation tool (MACC-Test)**

| **Methods** | **LST-LGA** | **LST-LPA** | **UBO-detector** | **Samseg** | **nnU-Net*** | **SynthSeg** | **WMH-DualTasker** |
| --- | --- | --- | --- | --- | --- | --- | --- |
| **Regions** |  | | | | | | |
| *frontal lobe (left)* | ${0.418}_{\pm0.110}$ | ${0.560}_{\pm0.088}$ | ${0.348}_{\pm0.110}$ | ${0.445}_{\pm0.092}$ | ${\mathbf{0.6}\boldsymbol{15}}_{\pm0.076}$ | ${0.505}_{\pm0.087}$ | ${0.602}_{\pm0.080}$ |
| *parietal-occipital lobe (left)* | ${0.375}_{\pm0.112}$ | ${0.535}_{\pm0.090}$ | ${0.338}_{\pm0.115}$ | ${0.420}_{\pm0.095}$ | ${\mathbf{0.6}\boldsymbol{08}}_{\pm0.078}$ | ${0.498}_{\pm0.090}$ | ${0.594}_{\pm0.083}$ |
| *temporal lobe(left)* | ${0.356}_{\pm0.115}$ | ${0.512}_{\pm0.092}$ | ${0.312}_{\pm0.096}$ | ${0.405}_{\pm0.109}$ | ${\mathbf{0.5}\boldsymbol{83}}_{\pm0.080}$ | ${0.472}_{\pm0.092}$ | ${0.518}_{\pm0.085}$ |
| *infratentorial region (left)* | ${0.345}_{\pm0.120}$ | ${0.510}_{\pm0.098}$ | ${0.322}_{\pm0.125}$ | ${0.388}_{\pm0.107}$ | ${\mathbf{0.5}\boldsymbol{78}}_{\pm0.083}$ | ${0.460}_{\pm0.095}$ | ${0.526}_{\pm0.221}$ |
| *basal ganglia (left)* | ${0.420}_{\pm0.108}$ | ${0.551}_{\pm0.088}$ | ${0.368}_{\pm0.112}$ | ${0.450}_{\pm0.090}$ | ${\mathbf{0.6}\boldsymbol{30}}_{\pm0.070}$ | ${0.518}_{\pm0.081}$ | ${0.620}_{\pm0.074}$ |
| *frontal lobe (right)* | ${0.405}_{\pm0.110}$ | ${0.570}_{\pm0.092}$ | ${0.353}_{\pm0.114}$ | ${0.440}_{\pm0.094}$ | ${\mathbf{0.62}\boldsymbol{2}}_{\pm0.074}$ | ${0.512}_{\pm0.086}$ | ${0.535}_{\pm0.079}$ |
| *parietal-occipital lobe (right)* | ${0.378}_{\pm0.112}$ | ${0.538}_{\pm0.094}$ | ${0.343}_{\pm0.117}$ | ${0.425}_{\pm0.095}$ | $\mathbf{0.612}_{\pm0.077}$ | ${0.502}_{\pm0.088}$ | ${0.496}_{\pm0.082}$ |
| *temporal lobe(right)* | ${0.353}_{\pm0.115}$ | ${0.505}_{\pm0.098}$ | ${0.315}_{\pm0.123}$ | ${0.398}_{\pm0.108}$ | ${\mathbf{0.5}\boldsymbol{80}}_{\pm0.082}$ | ${0.475}_{\pm0.094}$ | ${0.570}_{\pm0.087}$ |
| *infratentorial region (right)* | ${0.348}_{\pm0.120}$ | ${0.502}_{\pm0.099}$ | ${0.318}_{\pm0.126}$ | ${0.390}_{\pm0.110}$ | ${\mathbf{0.5}\boldsymbol{75}}_{\pm0.084}$ | ${0.468}_{\pm0.132}$ | ${0.562}_{\pm0.090}$ |
| *basal ganglia (right)* | ${0.423}_{\pm0.108}$ | ${0.560}_{\pm0.088}$ | ${0.375}_{\pm0.116}$ | ${0.455}_{\pm0.080}$ | ${\mathbf{0.6}\boldsymbol{35}}_{\pm0.068}$ | ${0.520}_{\pm0.080}$ | ${0.622}_{\pm0.099}$ |

**Note:** Table S7 reports the Dice similarity coefficient for region-wise performance across different WMH segmentation tools on the MACC-Test dataset. For each region, the method with the best performance is highlighted in bold. These evaluations provide insight into how different methods perform in specific brain regions, highlighting variations in segmentation accuracy due to anatomical complexity and method design. Dice scores are presented as mean ± standard deviation.

Table S8 **Region-level evaluation across all WMH segmentation tool (SINGER)**

| **Methods** | **LST-LGA** | **LST-LPA** | **UBO-detector** | **Samseg** | **nnU-Net*** | **SynthSeg** | **WMH-DualTasker** |
| --- | --- | --- | --- | --- | --- | --- | --- |
| **Regions** |  | | | | | | |
| *frontal lobe (left)* | ${0.427}_{\pm0.115}$ | ${0.575}_{\pm0.095}$ | ${0.382}_{\pm0.120}$ | ${0.465}_{\pm0.092}$ | ${\mathbf{0.6}\boldsymbol{15}}_{\pm0.080}$ | ${0.540}_{\pm0.090}$ | ${0.610}_{\pm0.085}$ |
| *parietal-occipital lobe (left)* | ${0.400}_{\pm0.118}$ | ${0.558}_{\pm0.100}$ | ${0.365}_{\pm0.123}$ | ${0.448}_{\pm0.105}$ | $\mathbf{0.610}_{\pm0.082}$ | ${0.525}_{\pm0.092}$ | ${0.595}_{\pm0.087}$ |
| *temporal lobe(left)* | ${0.352}_{\pm0.122}$ | ${0.502}_{\pm0.108}$ | ${0.325}_{\pm0.125}$ | ${0.410}_{\pm0.112}$ | ${\mathbf{0.5}\boldsymbol{88}}_{\pm0.088}$ | ${0.485}_{\pm0.096}$ | ${0.565}_{\pm0.090}$ |
| *infratentorial region (left)* | ${0.345}_{\pm0.130}$ | ${0.485}_{\pm0.112}$ | ${0.315}_{\pm0.128}$ | ${0.395}_{\pm0.115}$ | ${0.575}_{\pm0.090}$ | ${0.472}_{\pm0.097}$ | $\mathbf{0.590}_{\pm0.093}$ |
| *basal ganglia (left)* | ${0.450}_{\pm0.108}$ | ${0.600}_{\pm0.090}$ | ${0.400}_{\pm0.118}$ | ${0.485}_{\pm0.100}$ | ${0.650}_{\pm0.075}$ | ${0.560}_{\pm0.085}$ | ${\mathbf{0.66}\boldsymbol{0}}_{\pm0.084}$ |
| *frontal lobe (right)* | ${0.425}_{\pm0.115}$ | ${0.578}_{\pm0.095}$ | ${0.385}_{\pm0.120}$ | ${0.468}_{\pm0.102}$ | $\mathbf{0.628}_{\pm0.082}$ | ${0.542}_{\pm0.091}$ | ${0.612}_{\pm0.085}$ |
| *parietal-occipital lobe (right)* | ${0.402}_{\pm0.118}$ | ${0.560}_{\pm0.100}$ | ${0.368}_{\pm0.123}$ | ${0.450}_{\pm0.108}$ | $\mathbf{0.622}_{\pm0.064}$ | ${0.528}_{\pm0.092}$ | ${0.598}_{\pm0.087}$ |
| *temporal lobe(right)* | ${0.355}_{\pm0.122}$ | ${0.507}_{\pm0.158}$ | ${0.328}_{\pm0.125}$ | ${0.415}_{\pm0.112}$ | ${0.590}_{\pm0.088}$ | ${0.488}_{\pm0.096}$ | $\mathbf{0.608}_{\pm0.090}$ |
| *infratentorial region (right)* | ${0.348}_{\pm0.130}$ | ${0.488}_{\pm0.112}$ | ${0.313}_{\pm0.128}$ | ${0.398}_{\pm0.118}$ | ${0.578}_{\pm0.092}$ | ${0.475}_{\pm0.097}$ | $\mathbf{0.593}_{\pm0.092}$ |
| *basal ganglia (right)* | ${0.455}_{\pm0.118}$ | ${0.605}_{\pm0.082}$ | ${0.405}_{\pm0.126}$ | ${0.490}_{\pm0.088}$ | ${\mathbf{0.66}\boldsymbol{5}}_{\pm0.062}$ | ${0.565}_{\pm0.084}$ | ${0.635}_{\pm0.079}$ |

**Note:** Table S8 reports the Dice similarity coefficient for region-wise performance across different WMH segmentation tools on the SINGER dataset. For each region, the method with the best performance is highlighted in bold. These evaluations allow for a comparative understanding of method reliability across anatomical regions, with Dice scores reported as mean ± standard deviation.

Table S9 **Region-level evaluation across all WMH segmentation tool (MICCAI-WMH)**

| **Methods** | **LST-LGA** | **LST-LPA** | **UBO-detector** | **Samseg** | **nnU-Net**† | **SynthSeg** | **WMH-DualTasker** |
| --- | --- | --- | --- | --- | --- | --- | --- |
| **Regions** |  | | | | | | |
| *frontal lobe (left)* | ${0.445}_{\pm0.105}$ | ${0.610}_{\pm0.085}$ | ${0.410}_{\pm0.110}$ | ${0.500}_{\pm0.091}$ | NA | ${0.565}_{\pm0.083}$ | ${\mathbf{0.63}\boldsymbol{2}}_{\pm0.078}$ |
| *parietal-occipital lobe (left)* | ${0.428}_{\pm0.112}$ | ${0.590}_{\pm0.088}$ | ${0.395}_{\pm0.112}$ | ${0.485}_{\pm0.092}$ |  | ${0.551}_{\pm0.092}$ | ${\mathbf{0.62}\boldsymbol{4}}_{\pm0.081}$ |
| *temporal lobe(left)* | ${0.382}_{\pm0.115}$ | ${0.535}_{\pm0.095}$ | ${0.350}_{\pm0.116}$ | ${0.445}_{\pm0.098}$ |  | ${0.515}_{\pm0.092}$ | $\mathbf{0.588}_{\pm0.089}$ |
| *infratentorial region (left)* | ${0.375}_{\pm0.118}$ | ${0.520}_{\pm0.103}$ | ${0.345}_{\pm0.121}$ | ${0.435}_{\pm0.102}$ |  | ${0.508}_{\pm0.085}$ | $\mathbf{0.582}_{\pm0.087}$ |
| *basal ganglia (left)* | ${0.460}_{\pm0.102}$ | ${0.631}_{\pm0.082}$ | ${0.430}_{\pm0.113}$ | ${0.515}_{\pm0.092}$ |  | ${0.580}_{\pm0.082}$ | ${\mathbf{0.65}\boldsymbol{0}}_{\pm0.071}$ |
| *frontal lobe (right)* | ${0.450}_{\pm0.111}$ | ${0.615}_{\pm0.085}$ | ${0.415}_{\pm0.112}$ | ${0.505}_{\pm0.091}$ |  | ${0.568}_{\pm0.084}$ | $\mathbf{0.638}_{\pm0.078}$ |
| *parietal-occipital lobe (right)* | ${0.430}_{\pm0.111}$ | ${0.595}_{\pm0.088}$ | ${0.402}_{\pm0.112}$ | ${0.490}_{\pm0.093}$ |  | ${0.551}_{\pm0.085}$ | $\mathbf{0.625}_{\pm0.080}$ |
| *temporal lobe(right)* | ${0.385}_{\pm0.112}$ | ${0.540}_{\pm0.095}$ | ${0.355}_{\pm0.113}$ | ${0.448}_{\pm0.108}$ |  | ${0.518}_{\pm0.094}$ | ${\mathbf{0.59}\boldsymbol{0}}_{\pm0.087}$ |
| *infratentorial region (right)* | ${0.378}_{\pm0.120}$ | ${0.525}_{\pm0.099}$ | ${0.350}_{\pm0.126}$ | ${0.440}_{\pm0.110}$ |  | ${0.510}_{\pm0.132}$ | $\mathbf{0.585}_{\pm0.090}$ |
| *basal ganglia (right)* | ${0.465}_{\pm0.108}$ | ${0.635}_{\pm0.088}$ | ${0.435}_{\pm0.116}$ | ${0.525}_{\pm0.080}$ |  | ${0.582}_{\pm0.080}$ | $\mathbf{0.655}_{\pm0.091}$ |

**Note:** Table S9 reports the Dice similarity coefficient for region-wise performance across different WMH segmentation tools on the MICCAI-WMH dataset. For each region, the method with the best performance is highlighted in bold. These region-level evaluations highlight the segmentation accuracy of each method in anatomically distinct areas, with Dice scores reported as mean ± standard deviation. †nnU-Net is trained with the MICCAI-WMH dataset. Therefore, we cannot report the evaluation result on the same dataset.

Table S10 **Cluster-level evaluation across all WMH segmentation tool**

Cluster-wise evaluations assess the ability of each method to detect and segment individual lesion clusters. Metrics include Recall and F1-score measured at the cluster level. Cluster-wise Recall (↑) indicates the proportion of true lesion clusters correctly identified by the method, where higher values represent better performance. Cluster-wise F1 (↑) balances both precision and recall at the cluster level, reflecting the method’s overall accuracy in identifying and segmenting lesion clusters. Methods with the best performance for each metric are highlighted in bold. Results are presented as mean ± standard deviation, calculated from cross-validation experiments. †nnU-Net: Results on the MICCAI-WMH dataset are not reported, as the method was trained using this dataset. *p < 0.05, **p < 0.01, indicating statistically significant differences compared to WMH-DualTasker, as determined by the Wilcoxon signed-rank test with Bonferroni correction.

| **Dataset** | | | **MACC-Test (40)** | | **SINGER (64)** | | **MICCAI-WMH (60)** | |
| --- | --- | --- | --- | --- | --- | --- | --- | --- |
| **Methods** | **Input Modalities** | **Mask** | **Cluster-wise Recall** $\boldsymbol{\uparrow}$ | **Cluster-wise F1** $\boldsymbol{\uparrow}$ | **Cluster-wise Recall** $\boldsymbol{\uparrow}$ | **Cluster-wise F1** $\boldsymbol{\uparrow}$ | **Cluster-wise Recall** $\boldsymbol{\uparrow}$ | **Cluster-wise F1** $\boldsymbol{\uparrow}$ |
| **LST-LGA** | T1, T2-FLAIR | Yes | ${0.461}_{\pm0.110}$** | ${0.370}_{\pm0.085}$** | ${0.501}_{\pm0.076}$** | ${0.400}_{\pm0.064}$** | ${0.530}_{\pm0.053}$** | ${0.443}_{\pm0.073}$** |
| **LST-LPA** | T2-FLAIR |  | ${0.626}_{\pm0.090}$ | ${0.542}_{\pm0.085}$* | ${0.671}_{\pm0.056}$* | ${0.580}_{\pm0.055}$* | ${0.700}_{\pm0.045}$* | ${0.613}_{\pm0.063}$* |
| **UBO-detector** | T1, T2-FLAIR |  | ${0.400}_{\pm0.120}$** | ${0.311}_{\pm0.100}$** | ${0.363}_{\pm0.088}$** | ${0.362}_{\pm0.085}$** | ${0.478}_{\pm0.070}$** | ${0.383}_{\pm0.083}$** |
| **Samseg** | T1, T2-FLAIR |  | ${0.544}_{\pm0.094}$** | ${0.462}_{\pm0.080}$** | ${0.508}_{\pm0.073}$** | ${0.506}_{\pm0.072}$** | ${0.623}_{\pm0.063}$** | ${0.533}_{\pm0.063}$** |
| **nnU-Net*** | T2-FLAIR |  | ${\mathbf{0.6}\boldsymbol{80}}_{\pm0.070}$ | ${\mathbf{0.6}\boldsymbol{00}}_{\pm0.067}$ | $\mathbf{0.640}_{\pm0.058}$ | $\mathbf{0.645}_{\pm0.056}$ | $\mathrm{NA}$ | |
| **SynthSeg** | T2-FLAIR |  | ${0.582}_{\pm0.087}$* | ${0.502}_{\pm0.074}$** | ${0.540}_{\pm0.065}$** | ${0.544}_{\pm0.063}$** | ${0.645}_{\pm0.057}$** | ${0.563}_{\pm0.049}$** |
| **WMH-DualTasker** | T2-FLAIR | No | ${0.662}_{\pm0.078}$ | ${0.582}_{\pm0.064}$ | ${0.632}_{\pm0.051}$ | ${0.638}_{\pm0.050}$ | $\mathbf{0.758}_{\pm0.042}$ | $\mathbf{0.677}_{\pm0.053}$ |
